# Glutamine and GLS1 are key metabolic checkpoints that Limit T cell Infiltration and Function in HNSCC

**DOI:** 10.64898/2026.09.24.753717

**Authors:** Sonja-Maria Decking, Ines Ugele, Katja Dettmer, Gabriel S. Horkovics-Kovats, Julian Kuenzel, Charlotte L. Zuur, Joleen Traets, Clarissa Korf, Julian Todt, Fabian Kellermeier, Leah Voelkl, Torunn Zierul, Antonia Hehemann, Florian Westphal, Hanna Wenzel, Sigrid Wittmann, Laura Stengel, Stefan Loipfinger, Christian Schmidl, Vladimir M. Milenkovic, Eva Paddenberg-Schubert, Agnes Schroeder, Anja K. Wege, James A. Hutchinson, Paloma Riquelme, Christina Bruss, Luisa Symeou, Tobias Ettl, Ioannis Michaelides, Markus Kapsreiter, Maximilian Rink, Florian Weber, Daniel Simon, Istvan Pap, James S. McKenzie, Yuchen Xiang, Christopher Bohr, Peter J. Oefner, Marina Kreutz, Zoltan Takats, Kathrin Renner

## Abstract

Head and neck squamous cell carcinoma (HNSCC) is characterized by poor survival and limited benefit from immune checkpoint blockade (ICB), underscoring the need to define targetable mechanisms of immune escape. In a prospective, patient-centered study, we investigated how tumor metabolic reprogramming impairs T cell function and drives ICB resistance. Targeted metabolomics identified two dominant and distinct tumor states: the Warburg phenotype and glutamine deprivation and revealed increased tumor-specific glutamine to glutamate turnover. Integrating spatial metabolic imaging with immunohistochemistry linked these states to immune-cold and immune-excluded tumor architectures. Metabolic states selectively reduced the expression of T cell chemokine receptors highly abundant and prognostic in HNSCC. Accordingly, T cell migration into tumor spheroids was reduced at lower glutamine concentrations. Interfering with glutamine metabolism by pharmacologic inhibition or genetic deletion of glutaminase-1 markedly increased tumor susceptibility to immune cell-mediated cytotoxicity in vitro and ex vivo and sensitized patient-derived tumor fragments (PDTFs) to ICB. This effect extended beyond glutamine restoration and involved disruption of glutathione-dependent ROS defense, impaired tumor proliferation, and increased MHC expression in tumor cells. Consistently, high expression of genes involved in glutamine and glutathione metabolism associated with reduced ICB response in patients. In contrast to HNSCC, other tumor entities including breast cancer show low glutaminase-1 expression. Accordingly, breast cancer PDTFs were not responsive to glutaminase-1 inhibition. Together, these data identify glutaminase-1 as a metabolic checkpoint driving immune escape in HNSCC, and support guided patient stratification for metabolism-targeted immunotherapy, similar to current precision approaches for mutation-targeted drugs.

## Introduction

Head and neck squamous cell carcinoma (HNSCC) is among the most prevalent malignancies. Major risk factors are smoking, alcohol consumption and HPV infection (1, 2). Despite a variety of treatment strategies, outcome is low, recurrence rates are high and both have hardly improved in recent years (3, 4, 2). Nevertheless, a positive association between tumor-infiltrating T cells and progression-free survival (PFS) as well as overall survival (OS) has been shown (5, 6). Notably, T cell infiltration, function and localization as well as density of tumor-infiltrating cytotoxic T cells are of decisive prognostic relevance (7, 8). However, determining factors remain elusive.

During the last decade it became evident, that the function of immune cells depends on their metabolic activity, which in turn is determined by the metabolic microenvironment they reside in (9, 10). The accelerated metabolism of tumor cells results in low levels of nutrients as glucose and amino acids and the accumulation of metabolic end products as lactate and an acidic pH (11). This harsh conditions lead to a strong suppression of T cells and natural killer (NK) cells (12–16), as those nutrients are also of critical importance for immune cell activity and in particular T cell function (17, 18). Moreover, the differentiation and activity of immunosuppressive cell populations such as regulatory T cells (Treg) and myeloid-derived suppressor cells (MDSC) are fostered (11). In HNSCC, increased expression of glycolysis-related transporters and enzymes has been associated with a significantly worse prognosis (19, 20) and radioresistance (21). Furthermore, glutaminase (GLS), which converts glutamine to glutamate, is highly expressed in primary and metastatic HNSCC tissue (22, 23). Elevated expression of the glutamine transporter SLC1A5 associates with poor OS (24). These findings hint at accelerated glucose and glutamine metabolism in HNSCC, which we hypothesize might impact T cell infiltration and function, consequently, patient survival and response to immune checkpoint blockade therapy (ICB).

In 2017, ICB was approved in the palliative (25, 26) and, more recently, neoadjuvant setting for HNSCC patients (27). Nevertheless, only a minority of patients has shown a therapeutic response to ICB and long-term survival (25, 26).

We propose that targeting tumor metabolism may enhance anti-tumor immunity when combined with neoadjuvant therapy or ICB. Achieving this requires a detailed understanding of patient-specific tumor metabolic phenotypes and their impact on the immune microenvironment, as well as the identification of metabolic vulnerabilities that can be selectively exploited without compromising immune cell function.

To this end, we conducted a patient-centered study encompassing more than 240 tumor specimens, to capture tissue complexity and inter-patient heterogeneity. The Warburg phenotype and low glutamine levels were identified as the predominant and distinct key metabolic phenotypes, each associated with reduced T cell fractions. Spatial metabolic imaging integrated with immunohistochemistry linked these states to the established immune phenotypes, “inflamed”, “cold”, and “immune-excluded” tumors. Mechanistically, metabolic constraints selectively limited chemokine receptor expression, and glutamine availability governed T cell migration. We further identified glutaminase-1 (GLS1) as a key metabolic vulnerability: its inhibition curtailed tumor proliferation, impaired ROS defense, and enhanced susceptibility to immune-mediated cytotoxicity and ICB. Consistently, ICB-refractory tumors exhibited elevated expression of glutamine metabolism genes, highlighting the clinical relevance of these findings.

## Materials and Methods

### Study design

This study aimed on delineating the interplay of the metabolic tumor microenvironment and anti-tumor immunity in HNSCC thereby identifying and treatment strategies to increase anti-tumor immune response and ICB efficacy. A prospective non-randomized study was designed including 245 male and female patients diagnosed with primary or recurrent HNSCC at the University Hospital Regensburg. Cohort characteristics are described in Supplementary Table 1. Details on study participants and patient enrollment are given in Supplementary Material & Methods.

Studies involving human material were approved by the local ethics comitee (University Hospital Regensburg; votes HNSCC-study: 12-101-0070, 20-1891-101; vote breast cancer samples: 22-3151-101; votes healthy donor immune cells: 13-101-0240, 13-101-0238). All participants provided written informed consent and the study was conducted in accordance with the ethical standards of the declaration of Helsinki.

### Sample collection and storage

#### Blood

Blood for immune cell isolation was collected in lithium heparin tubes (Sarstedt), immediately inverted and kept at room temperature until further processing. For serum, blood was collected in a serum tube (S-Monovette, Sarstedt), centrifuged at 2,500 g for 10 min at 10 °C and stored at -80 °C.

#### Tissue specimens

HNSCC tissue specimens were obtained during tumor surgery or diagnostic panendoscopy performed under general anesthesia. Tissue was harvested before blood flow cessation to preserve the metabolic phenotype. For mass spectrometry based metabolite analysis, a part of each specimen was immediately snap frozen and stored at -80 °C by scientific personnel present in the operating room. Another subsample was immediately transferred into RNAprotect Tissue Reagent (Qiagen), kept at 4 °C overnight and stored at -20 °C until RNA isolation. Remaining tissue was quickly transported to the lab on ice and further processed for extraction of interstitial fluid, isolation of immune cells or tumor fragment culture.

BC tissue specimens were obtained as punch biopsies during surgery at the time of surgical clip placement prior to therapy initiation. Biopsies were immediately transferred into RPMI 1640 medium (Thermo Fisher Scientific) supplemented with 5% AB serum, 2 mM L-glutamine, insulin/transferrin/selenium/pyruvate (1×; Thermo Fisher Scientific), penicillin (50 U/mL; Thermo Fisher Scientific), and streptomycin (50 µg/mL; Thermo Fisher Scientific), and subsequently transported to the laboratory for further processing.

#### Interstitial fluid

The interstitial fluid (ISF) was collected according to the protocol of Wiig et al. (28). In brief, blood on tissue was removed either with tweezers or dabbed very carefully with a paper towel. Tissue was weighed, placed in the center of a 4 x 4 cm filter paper with a pore size of 15 µm (CellMicroSieves N15S, Thermo Fisher Scientific) and transferred to a 1.5 mL tube ensuring that the tissue was completely enclosed by the filter paper without squeezing it. Centrifugation was performed at 400 g for 10 min at 4 °C. The filter paper with enclosed tissue was removed from the tube. The retrieved tissue sample and ISF were immediately shock frozen and stored at -80 °C. Times between individual steps were kept as short as possible, all tubes and the centrifuge were pre-cooled and sample handling was performed at 4°C.

### Immune cell isolation from blood and tissue samples

Immune cells were isolated from lithium heparin blood by ACK-lysis and tissue samples after digestion with collagenase and DNase I. Details are given in Supplementary Material & Methods.

### Human T cell isolation, stimulation and culture

Peripheral blood mononuclear cells were isolated by density gradient centrifugation over Ficoll/Hypaque as described and CD3^+^ T cells were isolated by magnetic bead separation (Miltenyi Biotec). T cells were stimulated using anti-CD3/CD28 Dynabeads (Thermo Fisher Scientific) in a bead-to-cell ratio of 1:1 and cultivated in the presence and absence of 2 mM L-glutamine and 10 µM Bis-2-(5-phenylacetamido-1,2,4-thiadiazol-2-yl)ethyl (BPTES, Sigma-Aldrich) as described (29). T cell medium (RPMI1640) was supplemented with 10 % AB-serum (Bavarian Red Cross), penicillin (50 U/mL) and streptomycin (50 µg/mL, Thermo Fisher Scientific), 1 mM sodium pyruvate (Thermo Fisher), 1 % non-essential amino-acids (Thermo Fisher), 0.4 % MEM Vitamins (Thermo Fisher) and 50 µM β-mercaptoethanol and 25 IU/mL IL-2 (Thermo Fisher).

### Cell lines and origin

PCI-15 tumor cells (kindly provided by Prof. Dr. Richard Bauer, Department of Oral and Maxillofacial Surgery, University Hospital Regensburg) were cultivated in RPMI1640 (Thermo Fisher Scientific) supplemented with 10 % FCS and 2 mM L-glutamine in a humidified atmosphere (5 % CO_2_, 95 % air) at 37 °C. PCI-15 GLS1^-/-^ cells were generated using CRISPR/Cas9 (Details in Supplementary Data) and kept in cell culture medium supplemented with 20 mM glutamate (Sigma-Aldrich). Culture was controlled for mycoplasma contamination on a regular basis using the MycoAlert Mycoplasma Detection Kit (Lonza, Cologne, Germany) according to manufacturer’s instructions.

### Analysis of metabolic dependencies of tumor cell lines

0.2 x 10^6^ or 0.05 x 10^6^ tumor cells/well (for analysis after 24 h and 72 h, respectively) were seeded in 4 mL cell culture medium in 6-well plates and allowed to adhere. Subsequently, medium was removed, cells were washed and fresh medium containing the respective treatments. Details can be found in Supplementary Material and Methods.

### Co-culture of PCI-15 tumor spheroids and human immune cells

PCI-15 spheroids were directly seeded at a concentration of 0.05 x 10^6^ cells/100 µL to poly-hema coated 96 well U-bottom plates or generated with the hanging drop method (of 0.05 x 10^6^ cells/30 µL). Three days after seeding, spheroids were thoroughly washed to remove residual glutamine, and re-seeded to poly-hema coated 96-well U-bottom plates in 100 µL final volume and subjected to the respective treatments. Co-cultures investigating the effect of BPTES were performed in T cell medium to guarantee coherence with T cell experiments and to prevent BPTES precipitation. Co-culture was started on the following day. Leukocytes were isolated from blood of healthy human donors and co-culture was performed as described before (29). 72 h after starting co-culture, non-infiltrated immune cells were washed away, and co-cultures were further processed for flow cytometry analyses, Live Cell Imaging or embedded for IHC and DESI-MRM-MSI. Details can be found in Supplementary Material and Methods.

### Metabolite profiling

#### Determination of central carbon metabolites and amino acids in tissue homogenates and interstitial fluid

Extraction and quantification of metabolites from tumor tissues and ISF was performed as recently described (29). Details in Supplemental Material and Methods.

Tracing analyses: For tracing analyses 0.2 x 10^6^ in PCI-15 cells were seeded into 6 wells plates. After adherence, medium was replaced by T cell medium supplemented with 2 mM 13C5-glutamine (Hycultec) and BPTES (Sigma-Aldrich). After 24 h, cells were washed with ice-cold PBS and lyzed in ice-cold 80 % methanol. For T cells, isolation and culture was performed as described below in medium supplemented with 2 mM 13C5,15N2-glutamine (Hycultec) and BPTES. After 24 h, cells were washed twice with PBS and immediately frozen in liquid nitrogen. Metabolite extraction and GC-MS were performed as described previously (30).

#### Spatial metabolic imaging of tissue and spheroid sections

Snap frozen tissue was embedded and sectioned in a hydrogel matrix inside the cryostat chamber (Leica CM3050S), which had been cooled to -40 °C to prevent tissue thawing and facilitate the embedding process (31). DESI-imaging with 1 MRM transition per metabolite was performed at 25 µm spatial resolution for Figure 2 and 4 in negative ion mode using a TQ-XS mass spectrometer coupled to a Prosolia DESI stage with a high-performance sprayer (Waters). Details including optimized transition parameters are given in Supplemental Material and Methods. For spheroid sections, the same setup as for tissue sections was used with slight changes defined in Supplemental Materials and Methods. The imaging was performed with a spatial resolution of 25 µm and a scan time of 10 Hz. The images were acquired using HDI1.7 and MassLynx 4.1(Waters). The images were processed using HDI1.7.

#### Co-localization analysis of H/E-stained and metabolic images

To investigate the correlation between metabolite abundances and the pathological status of tissue at the cellular level, co-localization analysis was performed. H/E-stained images of the tissue samples were first bicubically down-sampled to match the pixel resolution of the metabolic images. Non-rigid affine transformations were then iteratively applied to maximize the Mander’s coefficient (MOC), which quantifies spatial overlap between the registered images.

For visualization, spatial distributions of the spectral ratios between lactate and glucose, as well as glutamine and glutamate, were calculated. Zero-intensity values were replaced with the average metabolite intensity within the tissue region. The final overlay depicted metabolites as heatmaps within the tissue microenvironment. Additionally, histopathologically annotated regions of high tumor cell concentration were aligned, transferred, and highlighted in red to illustrate spatial heterogeneity.

#### 3D sample preprocessing and coregistration

Raw MS imaging files were converted to MZ5 format using ProteoWizard/msConvert (version 3.0.24066-8d99481) and imported into MATLAB 2024b using in-house developed code. Image coregistration was also performed in MATLAB using in-house code and various functions from the Image Processing Toolbox. Details in Supplementary Material and Methods.

### H/E-staining

After DESI-MRM-MSI, slices were subjected to H/E staining. Details in Supplementary Material and Methods.

### Immunohistochemistry and TUNEL staining

IHC and TUNEL staining was performed on cryosections of tumor tissue, embedded spheroids and co-cultures as prepared for DESI-MRM-MSI. Details in Supplementary Material and Methods.

### Preparation of RNA, Reverse Transcription, and Quantitative Real-Time PCR

Isolation of RNA, reverse transcription and quantitative PCR were performed as described (32). Quantitative Real-Time PCR was performed using SsoAdvanced Universal SYBR Green Supermix (BioRad) with the CFXDuet Real-Time PCR System (BioRad). Primer sequences in Supplementary Material and Methods.

### ELISA

Interferon (IFN)γ and TNF were measured in culture supernatants by ELISA. All ELISA measurements were carried out using the respective R&D Systems Duoset Kit (R&D Systems) according to manufacturer’s protocols.

### Cell counting

Cell counts were determined using the CASY Cell Counter (Casy® Modell TT, Omni Life Science).

### Ex vivo tumor fragment culture (PDTF)

The protocol was adapted from Voabil et al. (33). Briefly, the tissue was cut into 15-20 mg fragments and exposed to the respective treatments for 24 h. Fragments were processed into single-cell suspensions for flow cytometry analysis of embedded in hydrogel for IHC. Details in Supplementary Material and Methods.

### Flow cytometry

Flow cytometry was performed using the BD FACS Fortessa X20 or FACS Celesta (BD Biosciences). Data were analyzed using the FlowJo Software (V10.9.0; Tree Star). Details on surface and intracellular staining protocols and determination of cellular ROS levels in Supplementary Material and Methods.

### Western Blot Analysis

PCI-15 and macrophages were cultivated with and without 2 mM L-glutamine for 24 h, detached, washed with PBS, suspended in Cell Lysis buffer (Cell Signaling Technology), and stored at -80 °C. Details in Supplementary Material and Methods.

### Statistical analysis

Statistical parameters including the exact number of experiments, the definition of the center, dispersion and statistical significance are reported in the figures and figure legends. Statistical analysis was performed with the GraphPad Prism software (V10). P-values below 0.05 were considered as statistically significant. Correlations were computed using Spearman’s R correlation test (non-parametric). Data were tested for normality using the D’Agostino & Pearson and, for smaller sample sizes, the Shapiro-Wilk test. For comparison of two groups, Mann Whitney (non-parametric, unpaired), Wilcoxon’s matched pair signed rank test (non-parametric, paired), or the paired t-test (parametric, paired) were applied. For two-group comparison of normalized data, a one-sample Wilcoxon signed rank test (non-parametric) or a one-sample t-test (parametric) against a hypothetical value of one were employed. For multiple group comparison, Kruskal-Wallis (non-parametric, unpaired) or Friedman’s (non-parametric, paired) with post-hoc Dunn’s test, respectively, or a RM one-way ANOVA (parametric, paired) or ordinary one-way ANOVA (parametric, unpaired) with Dunnett’s multiple comparison test were used. Asterisks in figures denote statistical significance (*p<0.05, **p<0.01, ***p<0.001). To adjust *p*-values for multiple testing, the Benjamini-Hochberg procedure was applied to control the false discovery rate at 0.05.

#### Survival analysis

Survival analysis was performed by log-rank (Mantel-Cox) test and plotted as Kaplan Meier estimation curves using the GraphPad Prism Software. For survival analysis of the in-house cohort cutoff was 1^st^ January 2024. Survival analyses on TCGA data were performed using the UCSC Xena browser (34). Data were retrieved from the TCGA Head and Neck Cancer dataset comprising gene expression data of 566 samples, after exclusion of normal tissue and metastasis 519 and 481 (with and without p16 positive samples, respectively) primary tumor specimen remained.

#### TCGA HNSCC data set

Raw read count data (HTSeq counts) for TCGA HNSCC patient cohort were retrieved using the GDCquery function from the *TCGAbiolinks* package (v1.15.1) in R (v4.0.2). Patients positive for HPV and normal tissue samples were excluded from the analysis. Raw count data were pre-processed and normalized using the *DESeq2* package (v1.30.0), including variance stabilizing transformation, for downstream analysis. Pearson correlation analysis was performed to assess the relationship between gene signature scores and expression of individual genes. Unsupervised clustering was conducted using k-means partitioning. The signature scores were calculated with the *GSVA* package (with Gaussian kernel, v1.38.2). Immune related signatures were obtained from Danaher et al. (35) and MCP counter (36). The hypoxia related signature was assessed using the 99-gene signature developed by Winter et al. (37). The IFNγ 10-gene signature was obtained from Ayers et al. (38).

#### IMCISION data set

Raw count data for the IMCISION cohort were obtained from Vos et al. (39), and normalization was performed using the *DESeq2* package (v1.30.0). Details regarding data generation and data preprocessing are provided in the original publication. Genes in the corresponding cluster obtained from the analysis on the TCGA HNSCC patient cohort were used to calculate the signature scores with the *GSVA* package (with Gaussian kernel, v1.38.2).

#### Single cell RNAseq data analysis

Gene expression of MCT1 and MCT4 was analyzed using single-cell RNA-seq data from HNSCC patients. Preprocessed and normalized gene expression data were obtained from Puram et al. (40) (GSE103322) including 5,902 single cells with 23,686 genes from primary and metastatic tumor tissue of 18 HNSCC patients. The provided cell type annotations were used, classifying 2,215 cells as malignant and 3,687 as non-malignant. UMAPs were generated for dimensionality reduction and visualization using the RunUMAP function in Seurat v5.2.1 on the provided normalized data (41). Gene expression of chemokine receptors, GLS and GCLC was analyzed using single-cell RNA-seq data approaching the UCSC Cell Browser (42). Preprocessed and normalized gene expression data were obtained from Conde-Lopez et al. (43, 44) (GSE164690, GSE181919, GSE234933), integrating scRNA-seq data from 78 patients (274,911 cells) across multiple studies.

#### Gene expression analysis in a TCGA Pan-Cancer dataset

Gene expression of GLS1 and GCLC was analyzed in a Pan-Cancer TCGA dataset. Data were retrieved from the UCSC Xena browser (34). Only primary tumor samples were included in the analysis, except for melanoma, which has been analyzed for primary tumors and metastases separately.

### Data and material availability

Throughout the manuscript, data are presented as single data points, representing underlying data. All datasets generated and analyzed in this study are available from the corresponding authors on reasonable request.

## Results

Investigating the expression of genes involved in glucose and amino acid metabolism in correlation to anti-tumor immune response identified two metabolism related clusters. Cluster 1 exhibited a negative correlation between glucose and glutamine metabolism related proteins, including glutaminase (GLS1) and amino acid transporters (SLC38A1, SLC38A2, SLC7A8, SLC7A5), with the immune infiltrate, including Th1 and cytotoxic T cells as well as IFNγ response (Figure 1A). In contrast, cluster 2, which included genes belonging to the SLC16A transporter family, mainly transporting aromatic amino acids, demonstrated no or a slightly positive association with T cell response (Figure 1A).

**Figure 1:**
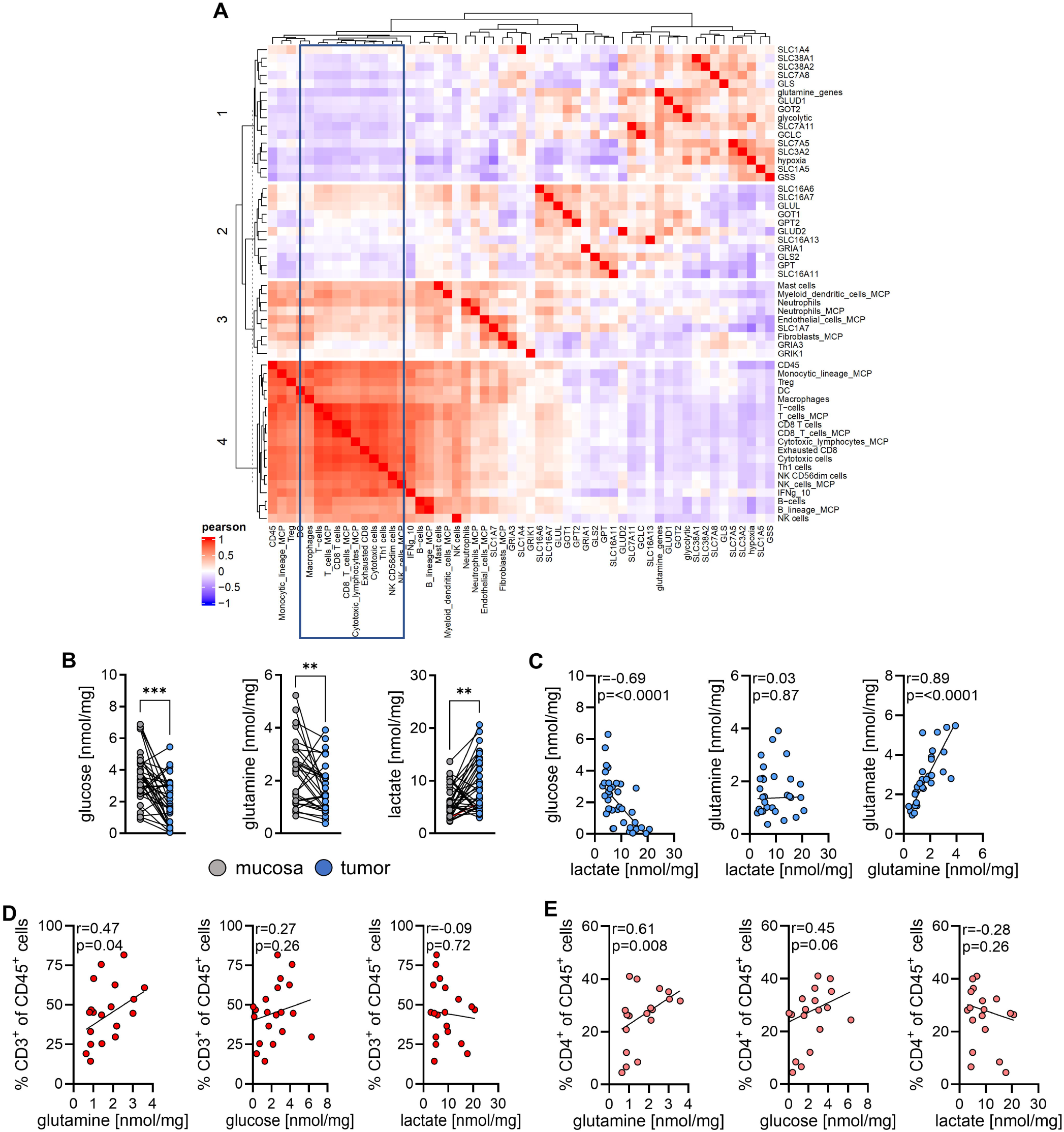
Metabolic alterations in HNSCC tumor tissue correlate with composition of tumor infiltrating immune cells. **(A)** Correlation plot (Pearson) generated with the HNSCC TCGA patient cohort between gene expression signatures, including immune infiltration related signatures and expression of individual genes related to glutamine, divided into four clusters (see Methods). **(B-E)** Tissue was harvested before blood supply cessation and immediately shock frozen in the operating theatre and stored at –80 °C. Metabolite levels were measured by mass-spectrometry. **(B)** Metabolite concentrations in homogenates of tumor tissue and matched mucosa. **(C)** Correlation analysis of selected metabolites extracted from tumor tissue homogenates. **(D, E)** Tissue was harvested and immediately processed. Single cell suspensions were generated from tissue and erythrocytes were removed. Immune cells were stained for population-specific surface markers and analyzed by flow cytometry. Correlations of **(D)** CD3^+^ T cells and **(E)** CD4^+^ T cell subpopulation with metabolite abundance are depicted. **(B, C)** Single values are shown matched between tumor and mucosa. Significance was calculated using a paired t-test for normally distributed data or otherwise the Wilcoxon matched-pairs signed rank test (**p<0.01, ***p<0.001). **(D, E)** Correlations were calculated using the Spearman r test.

We sought to translate these findings into phenotypic characteristics and to develop anti-metabolic therapeutic strategies capable of enhancing T cell responses and, ultimately, improving ICB efficacy. To this end a fully patient-centered study was initiated to cover inter-patient heterogeneity and the complex biology (patient characteristics Supplementary Table 1). To minimize effects of withdrawing blood supply, samples were taken before blood supply cessation. Clinical and medical scientists collaborated in the operating theatre to ensure immediate flash freezing of biopsies (within a minute) and rapid transfer (a five-minute walk) to the laboratory for further processing.

### Metabolic phenotypes correlate with intratumoral immune composition

To align the TCGA data set with our patient cohort, gene expression levels encoding respective metabolism related enzymes and transporters were analyzed (Supplementary Figure 1A). Tumor biopsies exhibited significantly elevated expression of amino acid transporters S*LC7A8*, glutaminase (*GLS1)* and glycolysis related genes such as lactate dehydrogenase A (*LDHA*), glucose transporter 1 (*GLUT1*) and monocarboxylate transporter (*MCT*) *1,* whereas *MCT4* was not increased, relative to matched mucosa (Supplementary Figure 1A). In accordance, targeted metabolomics revealed significantly reduced levels of glucose and glutamine and elevated lactate concentrations in tumor tissue (Figure 1B). Notably, other glycolytic and TCA intermediates did not differ in abundance, except for reduced citrate concentrations (Supplementary Table 2, Supplementary Figure 1B). It is noteworthy, that glutamine was the only amino acid significantly reduced in tumor tissue compared to mucosa. All other amino acids were either more abundant, as observed for proline (Supplementary Figure 1B), or did not differ discernibly (Supplementary Table 2).

Anatomic localization, staging and relapse status did not significantly impact metabolite levels, therefore, we abstained from stratification in subsequent analyses (Supplementary Figure 1C, data for relapse not shown).

The observed metabolic alterations indicate increased glucose metabolism and glutamine limitation as metabolic hallmarks. To further resolve metabolic heterogeneity, individual patient metabolite levels were pairwise correlated. Glucose and lactate showed a strong inverse correlation (Figure 1C), consistent with the Warburg phenotype in a subset of tumors. However, neither glucose (r=0.26, p=0.16; data not shown) nor lactate levels were related to glutamine concentrations (Figure 1C), whereas glutamine and glutamate showed a strong positive correlation (Figure 1C). Stratification of samples based on metabolite levels revealed distinct metabolic subgroups. Approximately 27 % of specimens showed no evidence of glucose or glutamine restriction (defined as below median) or lactate accumulation (defined as above median). In approximately 17 % of the samples the Warburg phenotype was detected, glutamine depletion alone was observed in 20 % and another 17 % displayed a mixed phenotype. Notably, lactate concentrations exceeding 10 mM, regarded as immunosuppressive (45, 46), were detected in 70 % of the “high lactate” samples. Finally, we quantified metabolite abundance in the interstitial fluid (ISF), the compartment immune cells reside in. The metabolic profile closely mirrored that observed in tissue (Supplementary Figure 1D; Supplementary Table 2). Metabolite correlations were likewise preserved (Supplementary Figure 1E). Notably, the lower glutamine to glutamate ratio in tumor-derived ISF in comparison to mucosa (0.53 vs. 1.35), indicates enhanced intra-tumoral conversion of glutamine to glutamate.

We next examined the relationship between metabolic phenotypes and leukocyte composition (gating strategy in Supplementary Figure 2A). Tumor tissue exhibited a marked reduction in CD3⁺ T cells, including both CD4⁺ and CD8⁺ subsets, relative to total leukocytes, accompanied by a concomitant expansion of the myeloid compartment (CD11b⁺), consistent across major myeloid populations (CD14⁺ and CD15⁺; Supplementary Figure 2B).

Correlation analyses revealed a significant positive association between CD3⁺ T cell frequency and glutamine levels, a weak with glucose and no association with lactate (Figure 1D). These relationships were more pronounced for the CD4^+^ T cell population (Figure 1E, Supplementary Figure 2C). Inverse correlations were observed between the CD11b^+^ myeloid compartment and glutamine (Supplementary Figure 2D) and CD15^+^ granulocytes and glucose and glutamine (Supplementary Figure 2E). In contrast the portion of CD14^+^ myeloid cells was not affected by metabolic conditions (Supplementary Figure 2F). In a small in-house cohort patients with a high proportion of CD4⁺ T cells (above median) exhibited a superior OS and PFS (Supplementary Figure 2G, Hazard Ratio 2.2 for OS and 2.6 for PFS).

### Intratumoral T cell distribution is related to metabolic phenotypes

Extraction of metabolites from homogenates results in loss of spatial and morphological information. Therefore, we subjected tumor sections to mass-spectrometry based metabolic imaging (MSI) and subsequent H/E staining, permitting the analysis of relative metabolite abundance and tissue morphology in the same section. We focused on metabolites of interest, glucose, lactate, glutamine and glutamate. To investigate whether single sections are representative, entire biopsies were processed. The metabolite pattern remained consistent throughout the entire specimen (Supplementary Figure 3A and 3B; 3D reconstruction in Supplementary Videos). Specifically in cancerous regions (annotation Supplementary Figure 4) MSI indicated the previously detected inverse abundance of glucose and lactate and the positive abundance between glutamine and glutamate (Figure 2A, Supplementary Figure 5A). Calculating spectral signal intensity ratios between lactate and glucose as well as glutamine and glutamate across whole tissue sections further supported this notion (Supplementary Figure 5B). To confirm regions of interest (ROIs) representative of tumor (T, yellow circle) and non-tumor tissue (NT, green circle) were defined (Supplementary Figure 5A), metabolite signal intensities were harmonized across samples (Supplementary Figure 5C) and extracted (Supplementary Table 3). The dynamic range of metabolite abundances obtained by absolute quantification (Figure 1B) and MSI signal intensities (Supplementary Table 3) was very similar (fold changes glutamine 11 vs. 10, glucose 136 vs. 114, glutamate 6 vs. 7, and lactate 10 vs. 5, respectively). The inverse relationship between glucose and lactate was recapitulated in T but not in NT areas (Figure 2B) and less pronounced in specimens displaying low lactate signal intensities (e.g. #12, #13, Supplementary Figure 5C). Conversely, glutamine and glutamate co-localized in both compartments; however, a steeper slope in tumor regions indicated again enhanced glutamine-to-glutamate conversion (Figure 2C).

**Figure 2:**
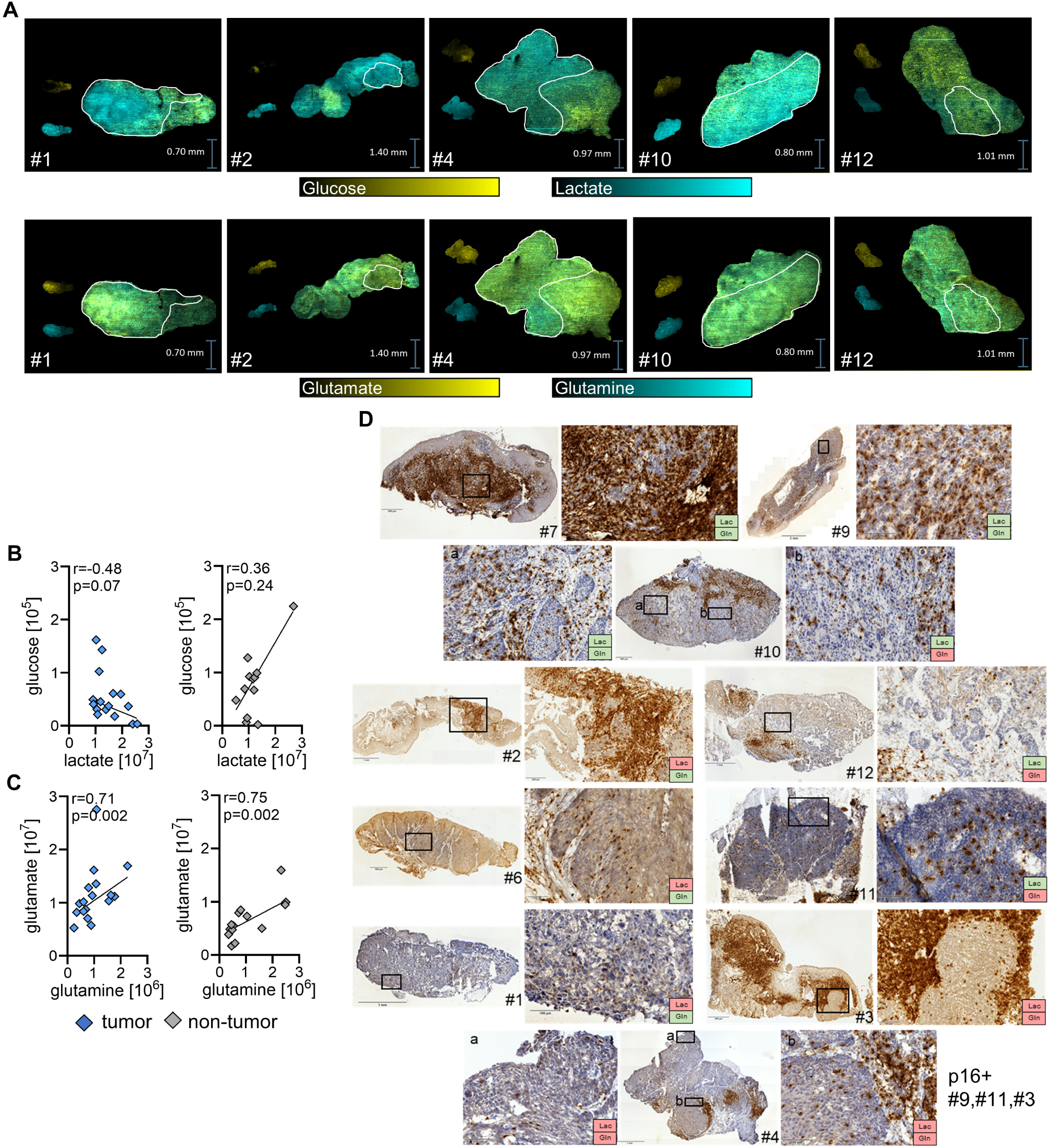
Metabolic alterations in HNSCC tumor tissue correlate with immune infiltration. Selected metabolites were analyzed with spatial resolution by DESI-MRM-MSI in 10 µm thick tumor tissue sections embedded in hydrogel. **(A)** Overlays of glucose (yellow) and lactate (blue) as well as glutamine (blue) and glutamate (yellow) are shown. Tumor cell dominated areas are encircled in white. **(B, C)** Regions of interest (ROIs, 70 scans, each scan 25 µm x 25 µm) representative for tumor (T) and non-tumor (NT) areas within a section were defined and mean signal intensities of respective metabolites were extracted (Table 2). Signal intensities of ROIs of paired tumor and non-tumor tissues and metabolite correlations for **(B)** glycolysis and **(C)** glutamine metabolism related metabolites are shown. Correlations were calculated using the Spearman r test. **(D)** CD3 expression was determined by immunohistochemistry in sections consecutive to those subjected to metabolic imaging. Samples were classified according to lactate (Lac) and glutamine (Gln) levels based on imaging signal intensities (Table 3). Low was classified as below median, high as above median signal intensity. Red Lac label: Lac high; Green Lac label: Lac low; Red Gln label: Gln low; Green Gln label: Gln high.

**Figure 3:**
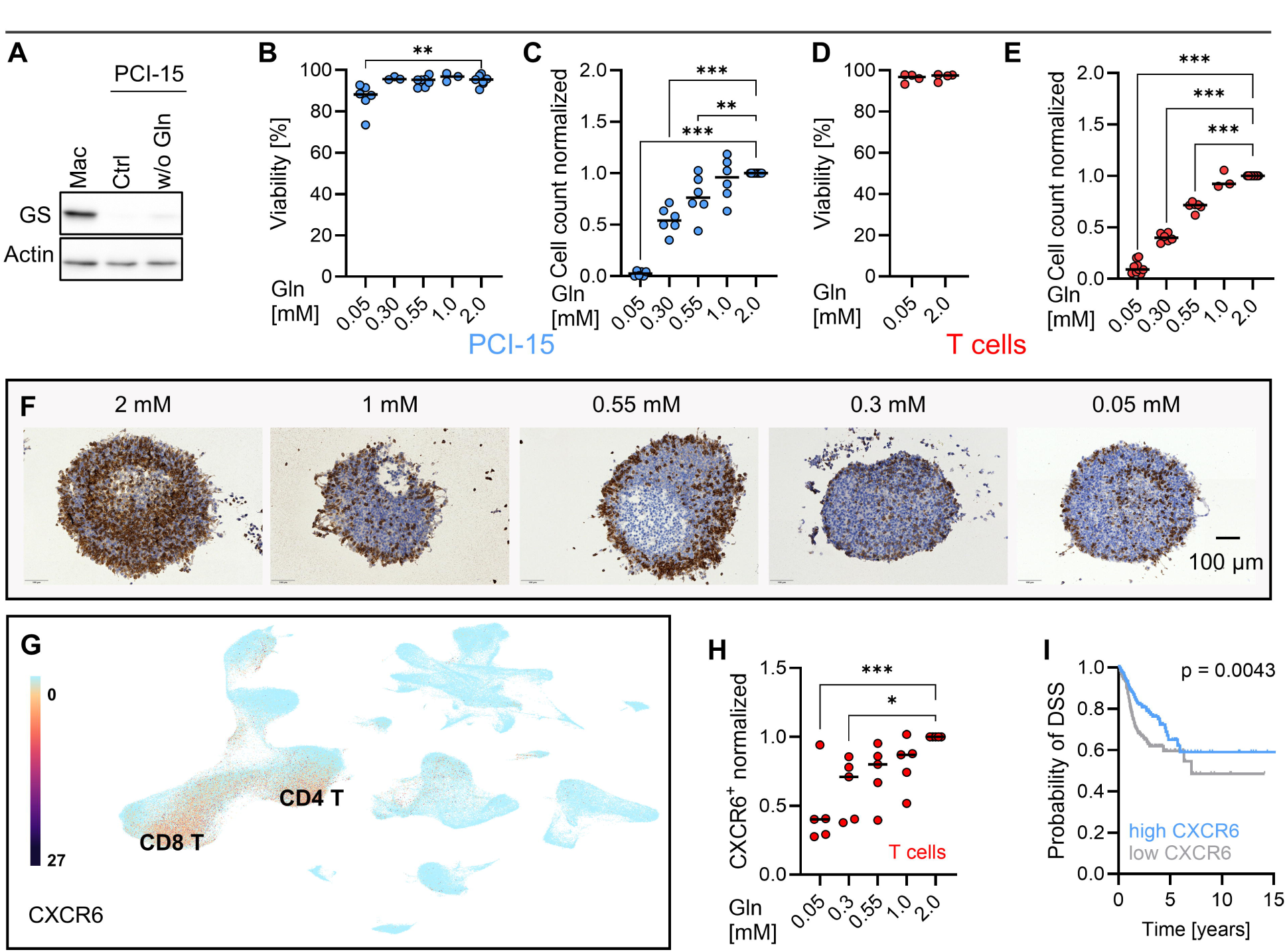
Glutamine restriction determines T cell infiltration and function. **(A)** Glutamine synthetase (GS) expression was analyzed by western blot, actin was used as loading control. Macrophages (Mac) served as positive control. **(B-F, H)** Cells were exposed to depicted glutamine (Gln) concentrations. Glutamine concentrations were calculated summing up medium concentration and 50 µM carry-over from serum supplementation. T cells were isolated from blood of healthy donors and stimulated with αCD3/CD28 Dynabeads. **(B)** Viability of PCI-15 cells was determined by flow cytometry analysis after Zombie NIR staining. **(C)** Cell count of PCI-15 cells was determined after 72 h. **(D)** Viability was determined by flow cytometry after Zombie NIR staining. **(E)** Cell count was determined after 6 days. **(F)** Co-culture of 3D spheroids with immune cells was performed. T cells were detected by IHC in spheroids. One representative experiment is shown (n = 3). **(G)** Expression of CXCR6 in HNSCC tumor and immune cells was analyzed in a publicly available single-cell RNAseq dataset approaching the UCSC cell browser. **(H)** CXCR6 expression of T cells was determined after 48 h by flow cytometry. **(I)** CXCR6 expression in relation to disease-specific survival (DSS) probability was analyzed in a TCGA dataset of Head and Neck cancer using the UCSC Xena browser. CXCR6 high was defined as above median expression. Survival was plotted as Kaplan Meier estimation curves. Significance was calculated applying the log-rank (Mantel-Cox) test, p<0.05 was considered significant.

T cells exhibit distinct spatial patterns within tumors, classified as inflamed (“hot”), immune-excluded (restricted to the stroma), and non-inflamed (“cold”) phenotypes, linked to clinical outcome [6]. We therefore analyzed spatial organization of T cell infiltration in sections adjacent to those used for MSI (Figure 2D). Samples were stratified according to metabolite signal intensities (Supplementary Table 2). Tumor specimens displaying signal intensity above and below median were subclassified into areas of high or low metabolite abundance. Tumors with pronounced T cell infiltration clustered mainly in low-lactate (green Lac label) and high-glutamine (green Gln label) groups, consistent with a favorable metabolic profile (Figure 2D).

In contrast, high-lactate tumors (red Lac label, Figure 2D) were predominantly cold or immune-excluded. Glutamine-low tumors (red Gln label) showed only limited T cell entry into tumor areas. Notably, intra-tumoral heterogeneity was evident in specimen #10, which contained both glutamine-high and glutamine-low regions; the latter showed reduced CD3⁺ T cell infiltration, further supporting a functional link between glutamine availability and T cell localization.

Retrospective annotation identified three p16⁺ tumors (#3, #9, #11), most likely HPV^+^, which were evenly distributed across metabolic and immune phenotypes. A small cohort of p16^+^ tumors showed metabolite distribution comparable to p16^-^ tumors (Supplementary Figure 6A). Including p16⁺ cases in survival analyses strengthened the association between CD4⁺ T cell abundance and clinical outcome (hazard ratios: 7.4 for OS and 3.7 for PFS; Supplementary Figure 6B). Accordingly, p16⁺ tumors were retained in all subsequent analyses.

### T cell infiltration depends on glutamine abundance

Our data indicate a link between metabolite abundance and T cell infiltration. While an inhibitory effect of lactate on T cell migration has been previously demonstrated (47, 48), the role of glutamine remains poorly defined. To address this, we employed our previously established 3D co-culture model of tumor spheroids with whole-blood leukocytes, which recapitulates key metabolic influences on T cell behavior in vivo (29). The HNSCC cell line PCI-15 was used as tumor model system. This cell line lacks glutamine synthetase (Figure 3A), which converts glutamate into glutamine, thereby enabling controlled manipulation of extracellular glutamine levels. Lactate was scarcely detectable in PCI-15 spheroids, excluding it as a confounding factor (data not shown). Glutamine availability did hardly impact viability (Figure 3B), whereas proliferation was significantly reduced at 0.55 mM glutamine or lower (Figure 3C). This was comparable to aCD3/CD28 stimulated T cells, which displayed preserved viability (Figure 3D) but proliferation was sensitive to glutamine limitation, with a significant reduction in the presence of a glutamine concentration (0.55 mM, Figure 3E), corresponding to the lower quartile of levels detected in tumor ISF. The expression of activation related markers, CD25 or CD226, was only affected by complete glutamine withdrawal (Supplementary Figure 7A and 7B). To avoid confounding effects, spheroids were generated in glutamine-containing medium, and T cells were pre-stimulated in the presence of glutamine before subsequent exposure to defined glutamine concentrations. Robust T cell infiltration also into the spheroid core were observed only at 2 mM glutamine, whereas at 1 mM and 0.55 mM T cell numbers were reduced and cells remained confined to the spheroid periphery (Figure 3F and Supplementary Figure 7C). To gain insights into underlying mechanisms, we assessed expression of chemokine receptors critical for T cell trafficking. Analysis of a publicly available single-cell HNSCC dataset (43) revealed CXCR6 and CXCR3 as predominantly expressed in tumor infiltrating T cells (Figure 3G and Supplementary Figure 7D-7F). Glutamine deprivation reduced CXCR6 but not CXCR3 expression (Figure 3H and Supplementary Figure 7G). Conversely, lactic acid lowered CXCR3 but not CXCR6 abundance (Supplementary Figure 7H). Both receptors strongly associated with disease-specific survival (Figure 3I and Supplementary Figure 7I). Collectively, these findings provide insight how metabolic constraints within the tumor microenvironment may limit T cell migration.

### Metabolic characteristics shape tissue resident T cell function

To investigate how metabolic states and potential anti-metabolic strategies affect tissue-resident T cells, we established an ex vivo patient-derived tumor fragment (PDTF) model. Immune profiles were compared in fresh biopsies (day 0) and after 24 hours of culture (day 1; gating in Supplementary Figure 2A and 8A). In some donors, relative T cell frequencies increased which was accompanied by a rather expected decline in CD15⁺ granulocytes (Supplementary Figure 8B). As we aimed to interfere with T cell function, IFNγ, perforin expression and portion of PD-1^+^ T cells (mean 42.88 ± 6.58 %) were assessed and remained unchanged (Supplementary Figure 8C), indicating preserved key functions.

To model Warburg tumors, fragments were cultured under glucose restriction or lactic acid exposure. Stepwise glucose reduction to complete withdrawal did not alter the percentage of IFNγ- or perforin-expressing T cells (Supplementary Figure 8D). While 10 mM lactic acid had no effect, 15 mM reduced IFNγ⁺ T cells, particularly in CD8⁺ cells (Supplementary Figure 8E). The relevance of IFNγ is underscored by its association with improved OS (Supplementary Figure 8F). Analysis of the Puram et al. single-cell HNSCC dataset (40) revealed higher MCT1 and MCT4 expression in tumor versus non-malignant tissue (Supplementary Figure 8G). Lactate secretion was markedly reduced in the presence of specific MCT1 and MCT4 inhibitors (Supplementary Figure 8H) and IFNγ^+^ and TNF^+^ T cells increased in three out of 5 donors (Supplementary Figure 8I). In responders, lactate reduction exceeded 50 %, whereas in a non-responder lactate declined only modestly (#3), suggesting ex vivo response might support patient stratification.

Glutamine dependency was examined by culturing fragments in 2.0 mM versus 0.55 mM glutamine (upper vs lower percentile in tumor tissue). Glutamine restriction immediately reduced IFNγ- and perforin-expressing T cells, particularly in specimens with high baseline production (Figure 4A). To interfere with glutamine metabolism, we focused on a clinically applicable GLS1 inhibitor guided by the following key observations. GLS1 expression was significantly elevated in tumor tissue (Supplementary Figure 1A), and tumors exhibited an increased glutamine to glutamate ratio. Furthermore, MSI data showed a strong co-localisation of glutamate with glutathione (Figure 4B), indicating that GLS1-driven conversion of glutamine to glutamate fuels glutathione synthesis and supports redox balance via ROS scavenging.

**Figure 4:**
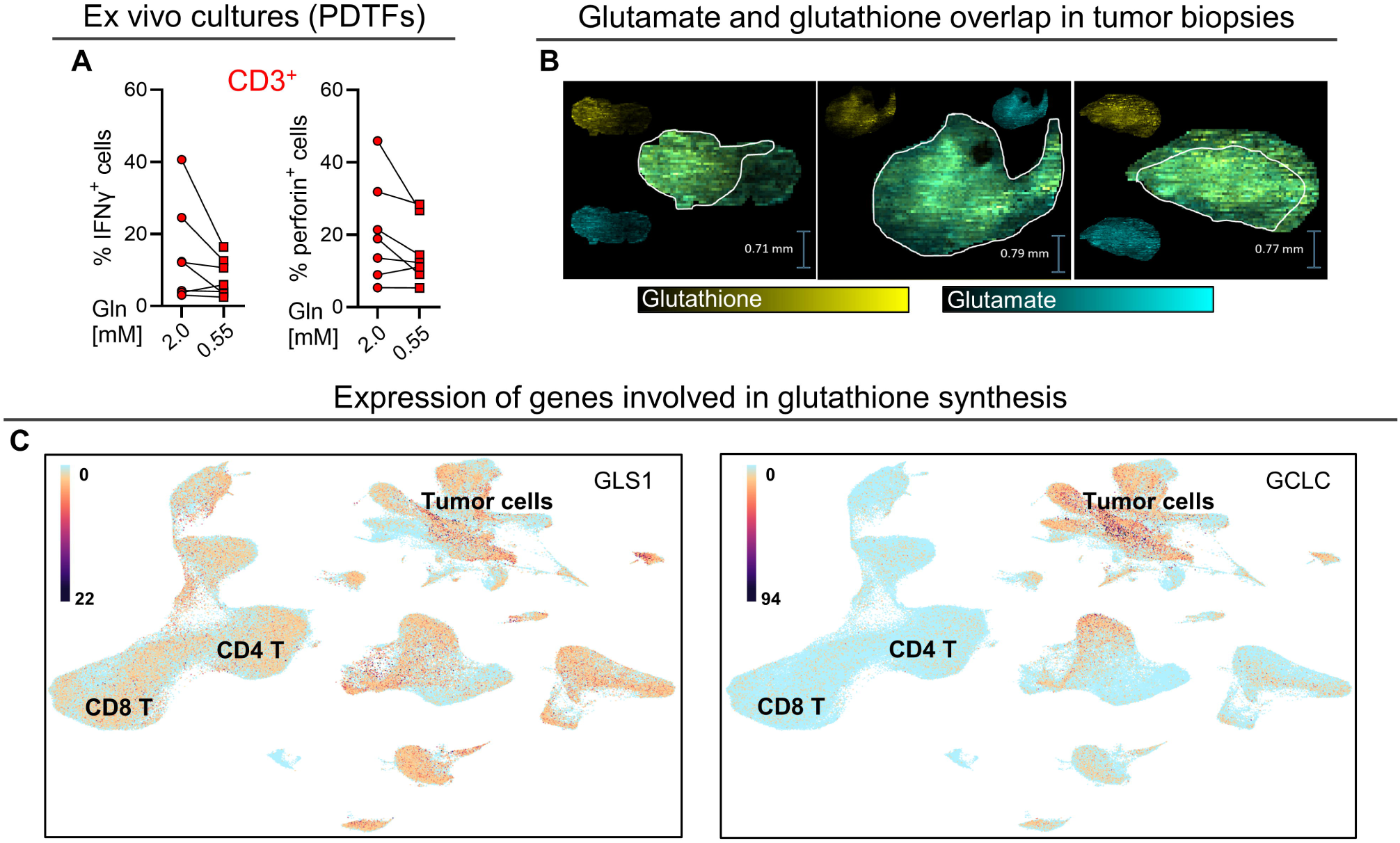
Relevance of glutathione synthesis in HNSCC. **(A)** Tissue fragments were subjected to respective glutamine concentrations for 24 h. Glutamine concentrations were calculated summing up medium concentration and 50 µM carry-over from serum supplementation. Single cell suspensions were prepared and proportion of IFNγ^+^ and perforin^+^ immune cell populations was investigated by flow cytometry. Single paired data points are shown. **(B)** Selected metabolites were analyzed with spatial resolution by DESI-MRM-MSI in 10 µm thick tumor tissue sections embedded in hydrogel. (**C**) Expression of GLS1 and GCLC in HNSCC tumor and immune cells was analyzed in a publicly available single-cell RNAseq dataset approaching the UCSC cell browser.

Investigating a single cell data set (44) revealed GLS1 expression in both T cells and tumor cells, irrespective of the HPV status (Figure 4C, HPV annotation depicted in Supplementary Figure 7E). In contrast, expression of the glutamate-cysteine ligase catalytic subunit (GCLC) was restricted to tumor cells, further supporting the particular relevance of glutathione for tumor cell redox homeostasis. The importance of ROS scavenging was underlined by the notion, that patients with a high expression of genes belonging to the ROS detoxification reactome exhibit a worse OS (Supplementary Figure 9A).

Given the limited data on human T cell function in the presence of a GLS1 inhibitor, we investigated effects in purified aCD3/CD28 stimulated T cell cultures and in our 3D co-culture model of tumor spheroids with immune cells, which predicted in vivo response to anti-metabolic treatment in our previous study (29). In both, PCI-15 and T cells, the conversion of glutamine to glutamate was strongly inhibited by BPTES (Supplementary Figure 9B), whereas intracellular glutamine peak areas were increased (Supplementary Figure 9C). In tumor cell culture supernatants extracellular glutamine concentration was elevated in the presence of BPTES (Supplementary Figure 9D). There was no discernible difference between 10 and 25 µM BPTES. Viability of T cells and PCI-15 cells was not affected by BPTES (Figure 5A). While proliferation of HNSCC cells was markedly impaired (50 %), T cell proliferation was only modestly reduced (10 %) in the presence of BPTES (Figure 5B). BPTES significantly increased ROS accumulation in PCI-15 cells, whereas ROS levels in T cells were hardly affected (Figure 5C). Consistently, glutathione levels were reduced in BPTES treated 3D PCI-15 spheroids (Figure 5D). Supplementation with glutamate or glutathione attenuated BPTES-induced ROS and proliferation defects in PCI-15 cells (Supplementary Figure 9E and 9F). BPTES did not alter IFNγ and TNF secretion (Figure 5E), in line with a report on sustained function of GLS1-deficient T cells (49). Glutamine restriction phenocopied BPTES treatment in PCI-15 cells, increasing ROS (Supplementary Figure 9G), impairing proliferation (Supplementary Figure 9H), and reducing glutathione in spheroids (Supplementary Figure 9I).

**Figure 5:**
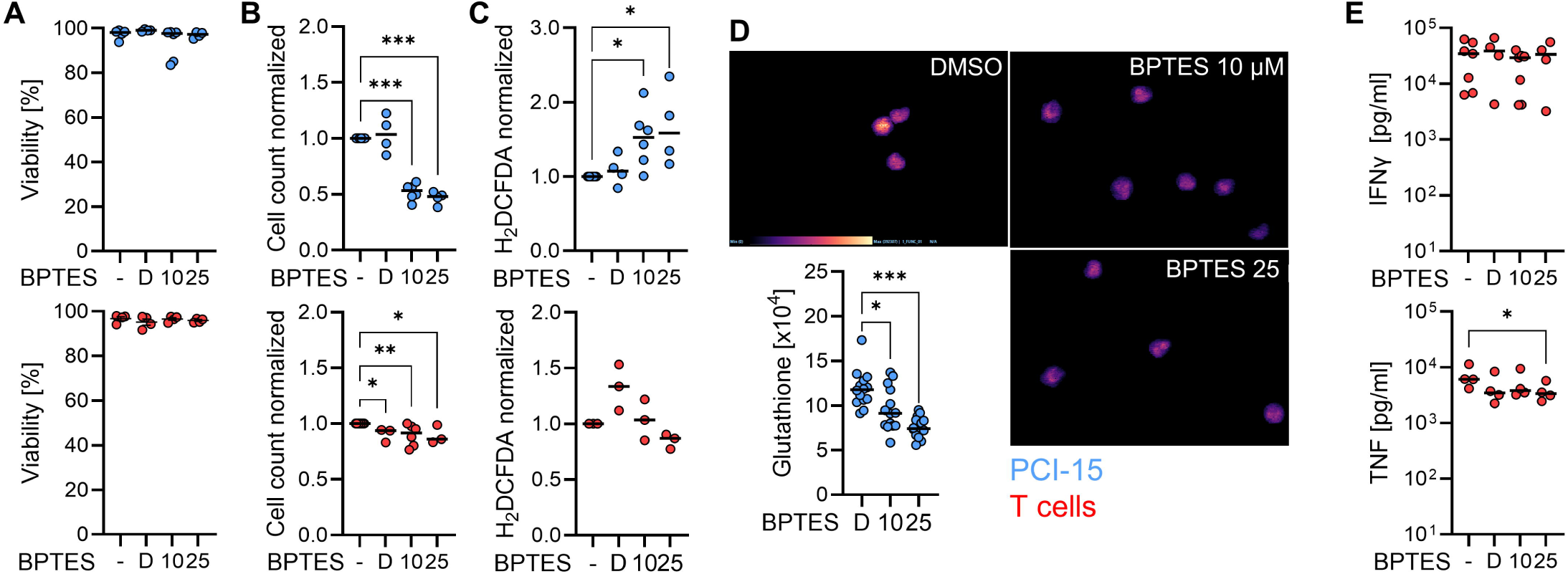
GLS1 inhibition reduces proliferation and disturbs redox balance in tumor cells without affecting T cell function. Cells were exposed to the indicated concentrations BPTES and the vehicle control DMSO. T cells were stimulated (αCD3/CD28 Dynabeads), PCI-15 cells were seeded and treated as indicated in medium with AB-serum. **(A)** Viability was determined by flow cytometry analysis. **(B)** Cells were counted after 72 h (PCI-15) and 6 d (T cells). **(C)** ROS production was assessed by H_2_DCFDA staining and flow cytometry analysis. **(D)** Spheroids were treated with the indicated BPTES concentrations and DMSO, embedded after 4 days and subjected to MSI for glutathione. Representative images (n=3) and extracted signal intensities from independent spheroids are displayed. **(E)** Cytokine concentrations in the culture supernatant of T cells were determined after 48 h. **(A – E)** Single data points and median levels are displayed. Significance was calculated using an ordinary one-way ANOVA and post-hoc Dunnett’s multiple comparison test, otherwise the Kruskal-Wallis test and post-hoc Dunnett’s multiple comparison test (*p<0.05, ** p<0.01, ***p<0.001).

In the absence of immune cells, BPTES did not affect spheroid viability (Figure 6A, w/o PBMCs). Immune cell dependent lysis, however, increased in the presence of 10 µM BPTES (Figure 6A – C, w/ PBMCs) and was not further enhanced at 25 µM (Figure 6C). To confirm specificity, GLS1 was knocked out in PCI-15 cells (GLS1⁻/⁻; Figure 6D). GLS1⁻/⁻ cells displayed reduced baseline proliferation, rescued by glutamate (Figure 6E), and were insensitive to BPTES (Figure 6F). GLS1⁻/⁻ spheroids exhibited enhanced immune-mediated lysis, mirroring BPTES-treated Mock spheroids, with no additional effect of BPTES (Figure 6G), thereby recapitulating and confirming GLS1-specific effects.

**Figure 6:**
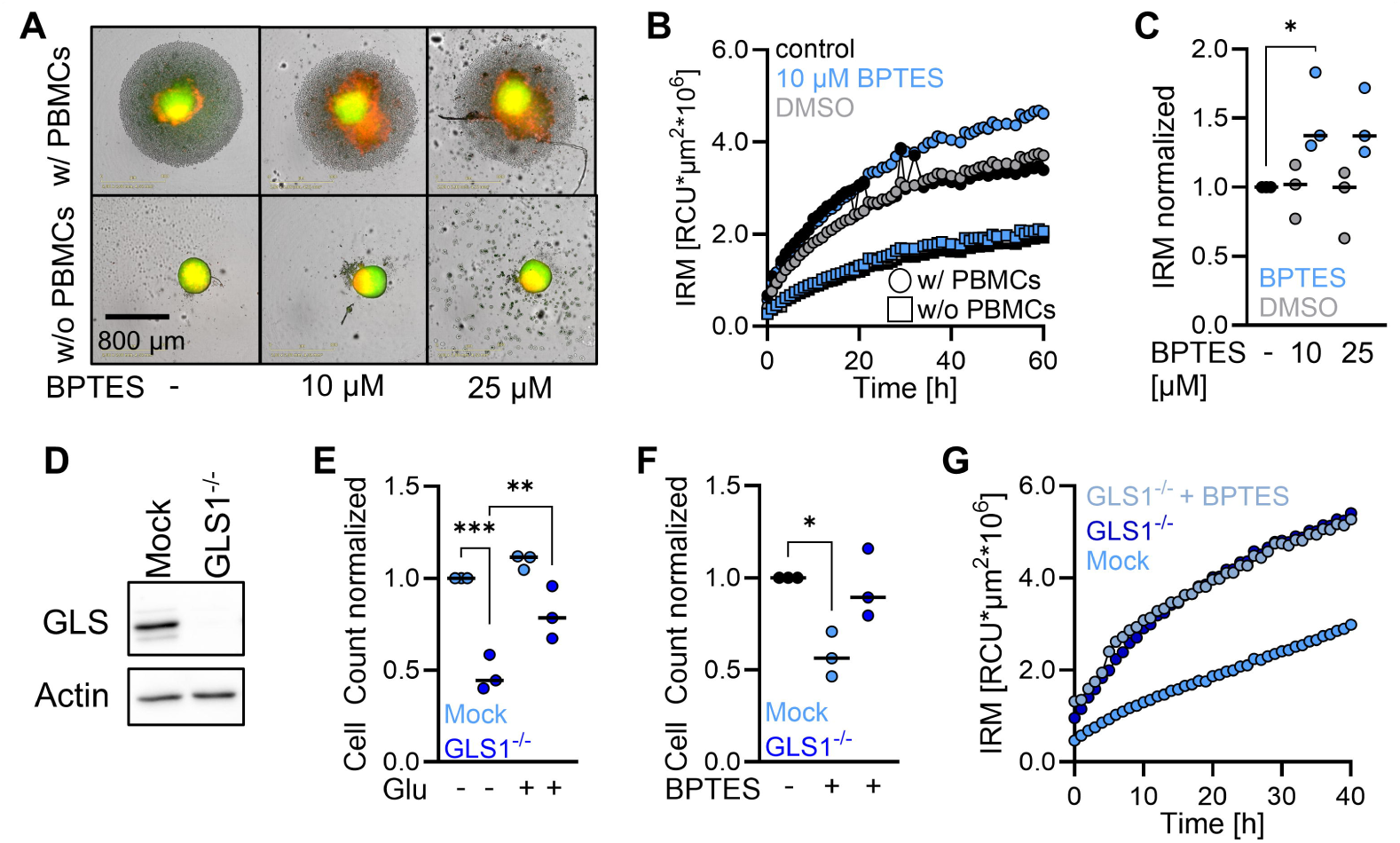
GLS1 inhibition enhances immune cell mediated spheroid lysis. **(A – C)** Spheroids were co-cultivated with (w/) and without (w/o) PBMCs and subjected to Live Cell Imaging. **(A)** One representative example is shown (n=3). Green color: living cells, red color: dead cells. **(B)** Integrated Red Mean Intensity (IRM) was analyzed over time (n=2). **(C)** Normalized IRM values after 60 h. **(D-G)** GLS1^-/-^ PCI-15 cells were generated by CRISPR/Cas9 technology. Cells treated with scrambled crRNAs (Mock) served as control. **(D)** GLS1 expression in Mock and GLS1^-/-^ cells was determined by Western blot. **(E, F)** Cells were cultivated **(E)** in the presence and absence of 20 mM glutamate (Glu) or **(F)** 10 µM BPTES and counted after 72 h. **(G)** Spheroids were co-cultivated with PBMCs in the presence and absence of 10 µM BPTES and subjected to Live Cell Imaging. IRM was analyzed over time (n=3).

Next, we evaluated BTPES in PDTFs. BPTES administration increased ROS in CD45^-^ cells and T cells (Figure 7A and Supplementary Figure 10A). Elevated ROS levels can be a sign of damage or T cell activation. BPTES did not reduce CD25, CD226, granzyme B or Ki-67 expression (Supplementary Figure 10B). Instead, it elevated the portion of IFNγ^+^ and TNF^+^ T cells (Figure 7B). The increase in IFNy^+^ T cells was reversed by glutamate or glutathione supplementation (Figure 7C). BPTES increased the portion of MHCI (HLA-ABC) expressing tumor cells (Figure 7D). To further characterize effects of GLS1 inhibition on tumor cells, we estimated apoptosis by determining DNA fragmentation in tumor-rich regions (H/E stains in Supplementary Figure 10C). BPTES consistently increased the percentage of apoptotic nuclei, with TUNEL^+^ nuclei predominantly within tumor areas (Figure 7E and Supplementary Figure 10C).

**Figure 7:**
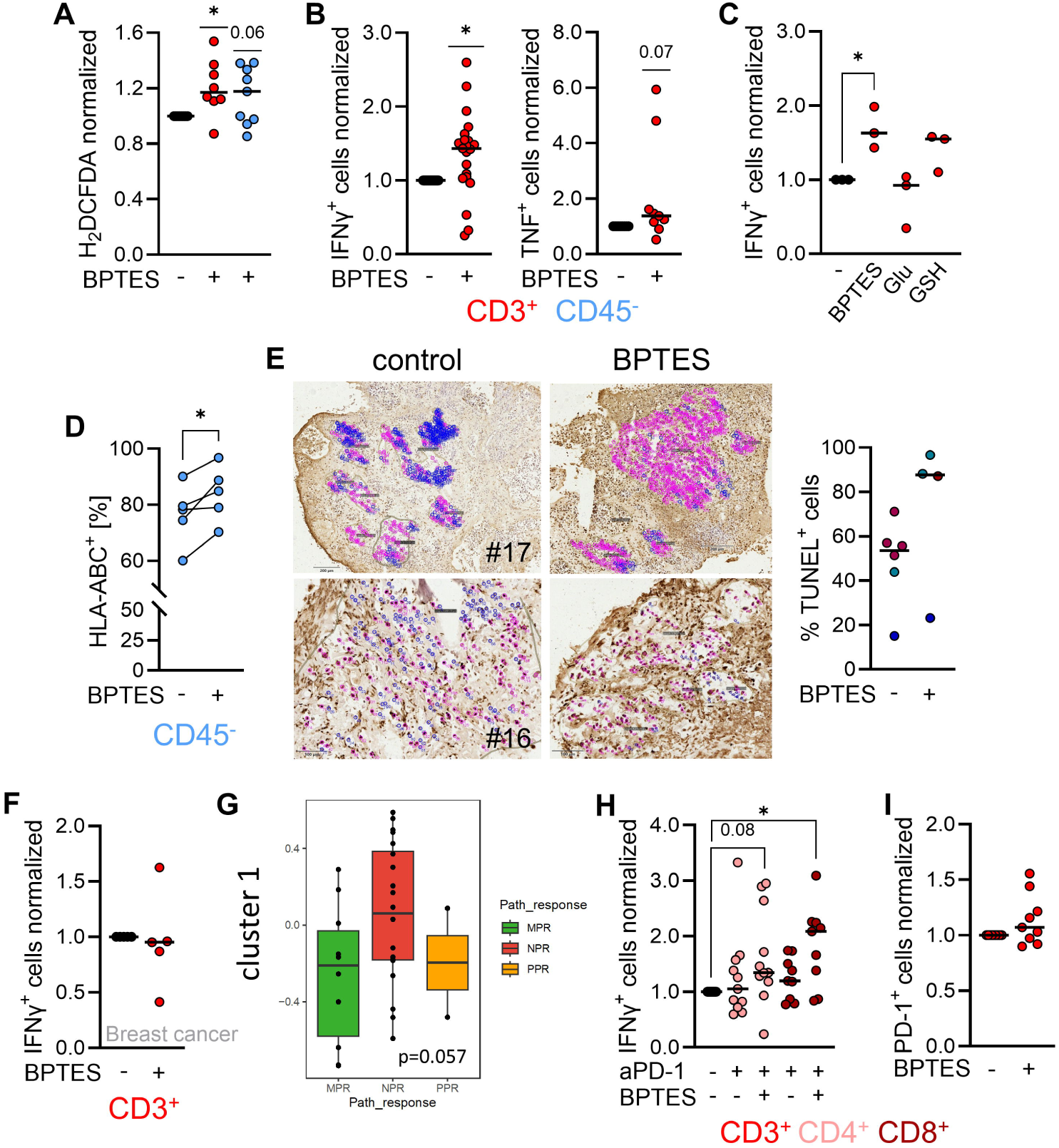
The GLS inhibitor BPTES increases anti-tumor immunity in patient-derived material. **(A, B)** Tumor fragments of HNSCC biopsies were subjected to 10 µM BPTES. After 24 h, single cell suspensions were prepared and analyzed by flow cytometry. **(A)** Intracellular ROS levels were investigated by H_2_DCFDA staining. **(B)** IFNγ and TNF expression in T cells was determined. **(C)** IFNγ expression in T cells was assessed in tumor fragments exposed to 10 µM BPTES with and without 20 mM glutamate (Glu) or 4 mM glutathione monoethyl ester (GSH). **(D)** HLA-ABC expression on CD45^-^ cells in tumor fragments exposed to 10 µM BPTES. **(E)** Tumor fragments were analyzed for apoptosis by TUNEL assay. Blue dots: TUNEL^-^ (viable), pink dots: TUNEL^+^ (dead) nuclei. Sections from two representative patients and summarized data of every counted region are depicted. Data of patients were marked with identical colors. **(F)** Tumor fragments of breast cancer biopsies were subjected to 10 µM BPTES. After 24 h, single cell suspensions were prepared and analyzed by flow cytometry for IFNγ expression. **(G)** Enrichment of gene cluster 1 (Figure 1B) in patients showing major pathological response (MPR) versus no pathological response (non-MPR) is shown. Statistical significance was assessed using the Wilcoxon rank-sum test. **(H)** IFNγ expression in T cells in fragments is shown, cultured with and without 10 µM BPTES and anti-PD-1 antibody (aPD-1, 10 µg/mL). **(I)** PD-1 expression was assessed in BPTES treated tumor fragments. (**A-F, H, I**) Single data points and median levels or paired data points are displayed. For two group comparison of normalized data, significance was calculated using the one-sample t test or the Wilcoxon signed rank test. For two-group comparison of raw values a paired t-test or the Wilcoxon test, for multiple group comparison the ANOVA and post-hoc Dunnett’s multiple comparison test was used (*p<0.05).

The GLS1 inhibitor Telaglenastat has been tested in clinical trials, however with limited efficacy. Comparative analysis of GLS1 expression across tumor entities revealed the highest expression in clear-cell renal cell carcinoma (ccRCC; KIRC) and the lowest expression in breast carcinoma (BRCA, Supplementary Figure 10D). Consistent with these findings, Telaglenastat showed limited efficacy in breast cancer (NCT03875313, NCT03057600), whereas ccRCC was the only tumor entity in which clinical trials reported promising outcomes (ENTRATA study, NCT03163667). HNSCC exhibited intermediate GLS1 expression but GCLC expression was among the highest in the analyzed tumor types, whereas BRCA again ranked among the lowest (Supplementary Figure 10E). To validate these observations and our results, we tested BPTES in PDTFs of breast cancer patients. In line with the clinical results, GLS1 inhibition failed to increase the frequency of IFNy^+^ T cells in breast cancer PDTFs (Figure 7F).

Finally, glutamine metabolism-high cluster 1 genes from the TCGA analysis (Figure 1B), which inversely correlated with T cell response, were enriched in ICB non-responders in the IMCISION trial (Figure 7G). This prompted combination of BPTES with anti-PD-1 (aPD-1) in PDTFs. In contrast to sole aPD-1 treatment, the co-administration of BPTES elevated significantly IFNγ-expressing CD4^+^ and CD8^+^ T cells (Figure 7H). BPTES neither affected the portion of PD-1 expressing T cells nor PD-L1 expressing myeloid or CD45^-^ cells (Figure 7I and Supplementary Figure 10F), indicating no direct checkpoint modulation.

Together, these data identify GLS1 as a critical metabolic checkpoint and support rapid bench-to-bedside translation of GLS1 inhibitors to enhance ICB efficacy in HNSCC.

## Discussion

Recent years have seen a paucity of effective treatment strategies for patients with HNSCC (50, 1). Although immunotherapy is now approved in both advanced and neoadjuvant settings, overall response rates remain modest (27). ICB efficacy critically depends on the presence and functional competence of tumor-infiltrating T and NK cells, yet the mechanisms regulating these responses are not fully understood. Accelerated and rewired tumor metabolism has emerged as a cancer hallmark that shapes the intra-tumoral immune landscape and impacts ICB response (51, 11, 15). Both, tumor cells and immune cells exhibit substantial and often competing demands for key nutrients such as glucose and glutamine (16). The heightened metabolic activity of tumor and stromal cells fosters a microenvironment characterized by nutrient depletion and the accumulation of metabolic intermediates and end products (17). Regarding HNSCC, elevated GLS expression (22, 23) upregulated glycolysis-associated transporters and enzymes (20, 19) correlate with poor outcome (24). However, in-depth characterization of metabolic phenotypes in conjunction with immune invasion phenotypes and function, as well as the evaluation of anti-metabolic treatment strategies remains lacking. To obtain reliable metabolic profiles, immediate snap-freezing of samples is mandatory, as metabolic adaptations to ischemia occur within seconds. The present study rigorously adheres to this critical requirement.

A comprehensive analysis of carbon metabolism in tumor and non-malignant tissue homogenates, interstitial fluid, and by spatial metabolic imaging identified elevated glycolysis, low glutamine concentrations, and increased glutamine-to-glutamate conversion as key, but partly independent, alterations in tumors. Glutamine was the only amino acid consistently reduced in tumor tissue versus matched mucosa. These metabolic features correlated negatively with T cell proportions among infiltrating leukocytes and with immune phenotypes (hot, cold, T cell–excluded), where cold and excluded tumors portend poor prognosis (7, 8, 6). In contrast, tumors with dense T cell infiltration, including in tumor cell–rich regions, clustered in a metabolically favorable group with low lactate and high glucose/glutamine levels. While lactate is known to impair T cell migration and infiltration (14, 47, 52), the role of glutamine in T cell trafficking remains less defined.

A role of the mTORC1 complex, often regarded as glutamine sensor, in coordinating cell migration has been described (53), and CD8^+^ T cell motility declines under glutamine deprivation (54). Consistent with this, we observed that low glutamine conditions confine T cells to the spheroid periphery and reduces overall presence. Mechanistically, high lactic acid and low glutamine reduced expression of the chemokine receptors CXCR3 and CXCR6, with lactic acid mainly affecting CXCR3 and glutamine availability regulating CXCR6, indicating selective metabolic control of chemokine receptors. CXCR3 and CXCR6 mediate T cell entry into tumors and hold strong prognostic value for outcome and ICB response across cancers, including HNSCC (55–60), yet direct metabolic regulation of their expression has been little explored. Our findings suggest that normalizing metabolic conditions can promote T cell infiltration, revealing a previously underappreciated link between tumor metabolism and immune cell trafficking.

Metabolic conditions not only shape T cell migration but also impair tissue-resident T cells. The functionality of immune cells is contingent upon their metabolic activity (9, 61) and enhanced tumor glycolysis and glutaminolysis are both immunosuppressive (30, 62, 14, 29, 63). Increased glycolytic flux with extracellular lactic acid accumulation diminishes T and NK cell function and viability (12, 13). Consequently, anti-glycolytic interventions could restore T cell functionality and ICB efficacy. In this study, combined MCT1 and MCT4 inhibition increased T cell function in specimens in which lactate secretion was reduced by approximately 50 %.

The pivotal role of glutamine for T cell function is well established (64, 65) and is reinforced by our data. However, human T cells exhibited no functional impairment in the presence of the GLS1 inhibitor BPTES, contrary to expectations from studies implicating a role for glutamine in T cell ROS defense (64). Our findings are consistent with a recent study reporting maintained anti-tumor function of T cells lacking GLS1 expression (49) and reports that GLS1 inhibition promotes Th1 differentiation and strenghtens anti-tumor response (66, 62). Here BPTES increased the frequency of IFNγ^+^ T cells and induced higher apoptosis rates in PDTF, a model system shown to be of clinical relevance (33). Further supporting this concept, in-depth analysis of data from the IMCISION trial revealed elevated glutaminolysis related genes in patients not responding to ICB. These findings provide the rationale for combining ICB with metabolic inhibitors, such as the GLS1 inhibitor CB-839 or the MCT1 inhibitor AZD3965 (NCT01791595). Encouraging results with such combination strategies have been observed in other cancer entities (67). The therapeutic potential of targeting glutamine metabolism in HNSCC is underlined by the synergistic antitumor activity observed with the GLS1 inhibitor CB-839 and CPI-613, a novel lipoate analog that promotes glutaminolysis (68). However, GLS1 inhibition enhanced ICB response in preclinical models (63, 69), clinical trials have shown more modest results. Even combination approaches, such as Telaglenastat with paclitaxel or checkpoint inhibitors, have yielded manageable safety profiles but only modest antitumor activity in most solid tumor settings. This gap between preclinical promise and clinical outcome likely reflects either compensatory metabolic rewiring or limited patient stratification by biomarkers. Our analysis showed low GLS1 expression in breast cancer, and BPTES treatment had no effect in breast cancer PDTFs, consistent with clinical trial outcomes. This demonstrates that careful patient stratification is essential for the successful implementation of anti-metabolic treatment strategies and should be taken into account when designing and conducting clinical trials.

## Acknowledgments

The PCI-15 cell line was kindly provided by Prof. Richard Bauer (Clinic and Polyclinic for Oral and Maxillofacial Surgery, University Hospital Regensburg). We thank Monika Wehrstein, Ute Schreiter, Claudia Wögerbauer and Sona Hakobyan from University Hospital Regensburg for excellent technical assistance. We thank the FACS Analytics and Cell Sorting Core Facility of the Leibniz Institute for Immunotherapy for supporting flow cytometry experiments.

## Funding

The study was financially supported by following institutions:

Wilhelm Sander Foundation, project number 2021.137.1 (KR, IU)

Else Kröner-Fresenius Foundation (IU)

Bayerisches Zentrum für Krebsforschung (BZKF, Bavarian Center for Cancer Research), lighthouse project “Pre-Clinical Models” (KR, SMD)

ReForM E program (IntraCom consortium) University Hospital Regensburg, Germany

## Author contributions

KR conceived the project, supervised experiments, analyzed data, and wrote the manuscript. IU and SMD performed and supervised experiments, analyzed data, and wrote the manuscript. CK, JT, LV, TZ, CL, LS, FWes, HW performed experiments and analyzed data. SW, LS, TE, IM, MKa, AKW have acquired patients for the study and taken sample material. JK and MR acquired patients and revised the manuscript. KD and FK performed metabolomics analyses. GSHK, AH, DS, IP, YX, JM performed DESI-MRI experiments. CZ, JT, CS and SL performed and supported bioinformatical analyses of transcriptomics data. VM assisted with methodological development. FWeb supported immunohistochemistry analyses. CB, PJO, MKr and ZT provided critical resources and revised the manuscript. EPS, AS, JAH and PR revised the manuscript.

