## Supplementary Data for "Glutamine and GLS1 are key metabolic checkpoints that Limit T cell Infiltration and Function in HNSCC"

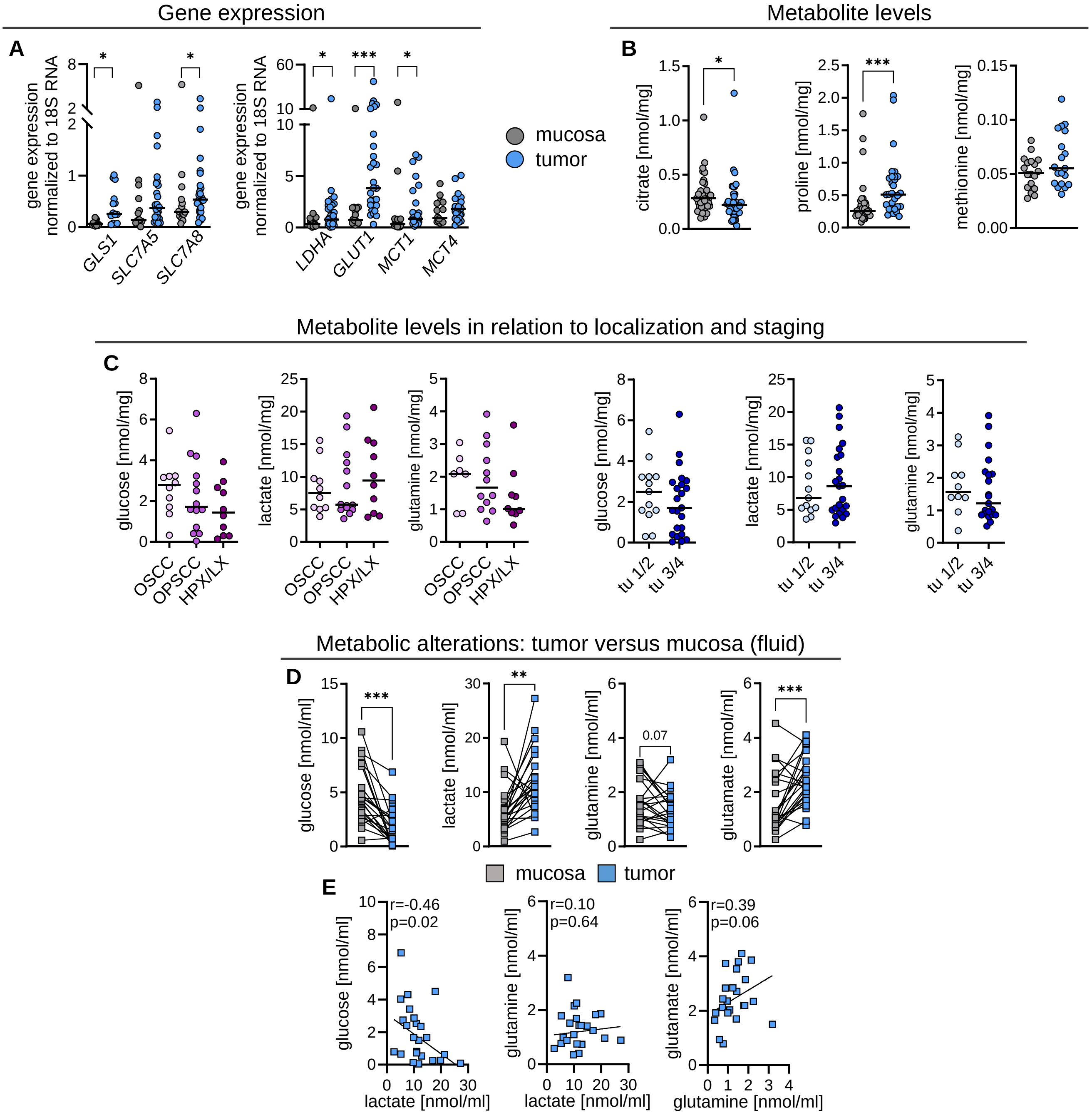


**Supplementary Figure 1. Tumor related alterations in expression of metabolism associated genes, metabolite levels and impact of localization and staging thereof.** (**A**) Tumor tissue and mucosa were harvested and immediately stored in RNAprotect. After RNA isolation, expression of respective genes was analyzed by RT-qPCR and normalized to 18S RNA expression. **(B, C)** Tissue was harvested and immediately shock frozen in the operating theatre and stored at –80 °C. (**B**) Metabolites were determined by mass spectrometry-based methods in homogenates of tumor tissue and matched mucosa. (**C**) Metabolite levels were analyzed in relation to tumor location (OSCC = oral squamous cell carcinoma, OPSCC = oropharyngeal squamous cell carcinoma, HPX/LX = hypopharynx/larynx) and staging as determined by tumor size (tu = tumor, 1/2 = tumor stage T1-2, 3/4 = tumor stage T3-4 according to UICC). **(D, E)** Tissue was removed and immediately transported to the laboratory on ice to retrieve interstitial fluid. **(D)** Metabolite concentrations in the interstitial fluid of tumor tissue and matched mucosa. **(E)** Correlation analysis of selected metabolites extracted from interstitial fluid of tumor tissue. **(A-E)** Shown are single values and median levels. For two group comparison significance was calculated using the Mann Whitney test, for multiple group comparison Kruskal Wallis and post-hoc Dunn's multiple comparison test was used (*p < 0.05, ***p < 0.001). Correlations were calculated using the Spearman r test.

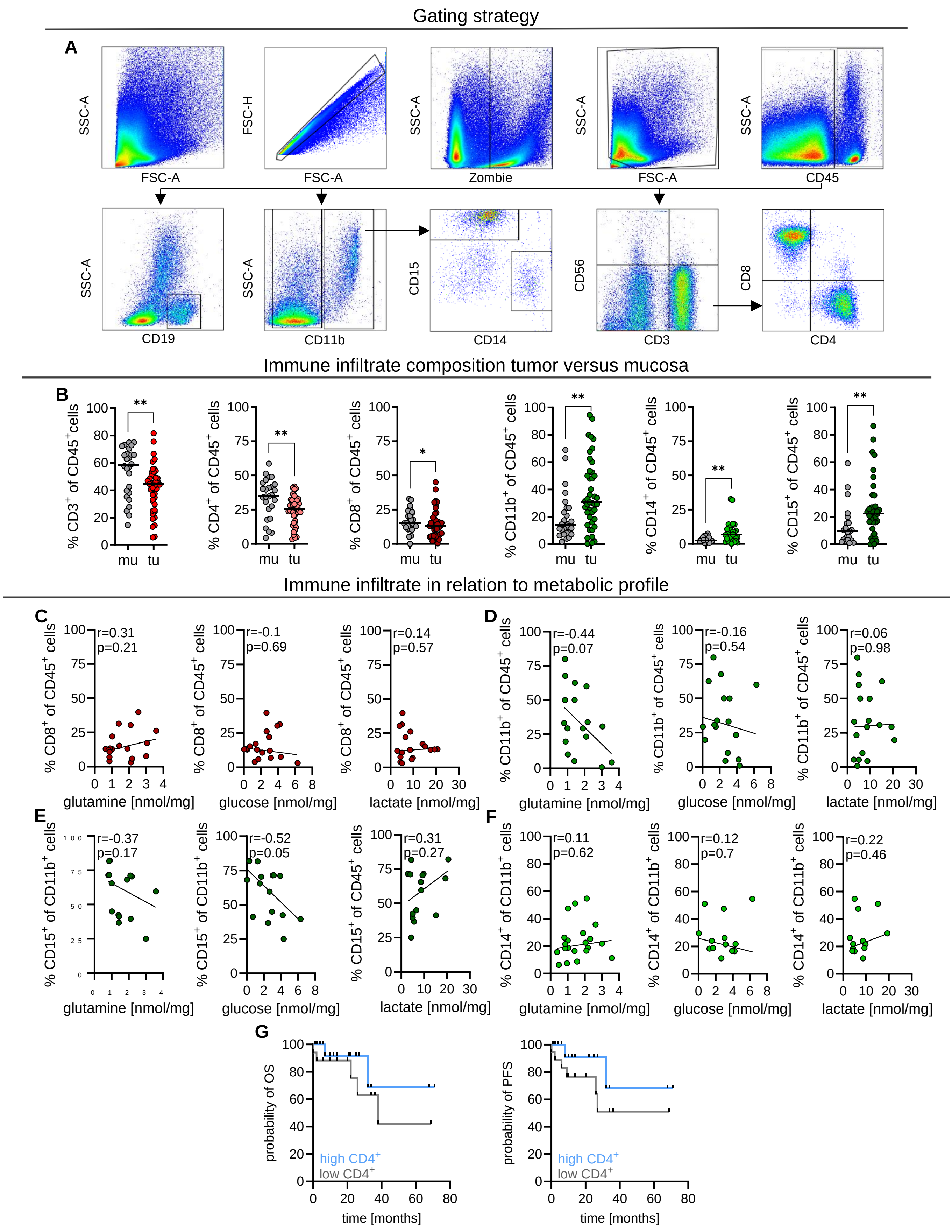


**Supplementary Figure 2. Alterations in immune cell composition in tumor tissue is related to metabolic features. (A-G)** Tissue was harvested and immediately processed. Single cell suspensions were generated from tissue and erythrocytes removed. Immune cells were stained for population-specific surface markers and analyzed by flow cytometry. (**A**) Gating strategy for one representative donor is displayed. (**B**) Specific immune cell populations among $CD45^+$ leucocytes isolated from mucosa (mu) and tumor (tu) tissue. Displayed are single values and median levels. Significance was calculated using the Mann Whitney test (*$p < 0.05$). Correlations of **(C)** $CD8^+$ T cell, **(D)** $CD11b^+$ myeloid cell proportion and respective (**E**) $CD14^+$ and (**F**) $CD15^+$ subsets with metabolite abundance are depicted. Shown are single data points and correlations were calculated using the Spearman R test. (**G**) Clinical data on overall and progression-free survival (OS/PFS) of patients were collected. Shown is OS and PFS dependent on the percentage of $CD4^+$ T cells. Survival was plotted as Kaplan Meier estimation curves.

3D metabolite distribution

**A**

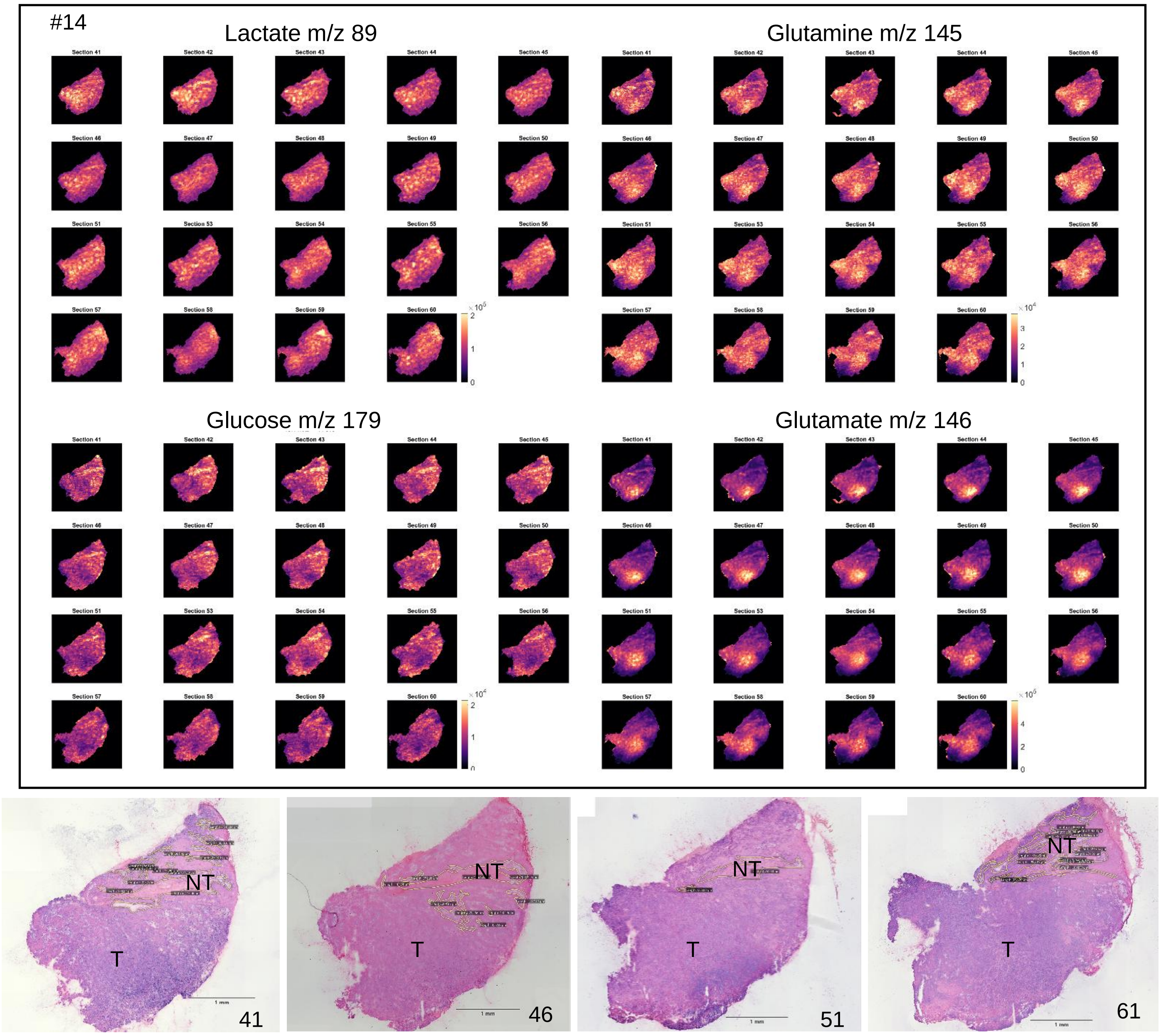

**B**

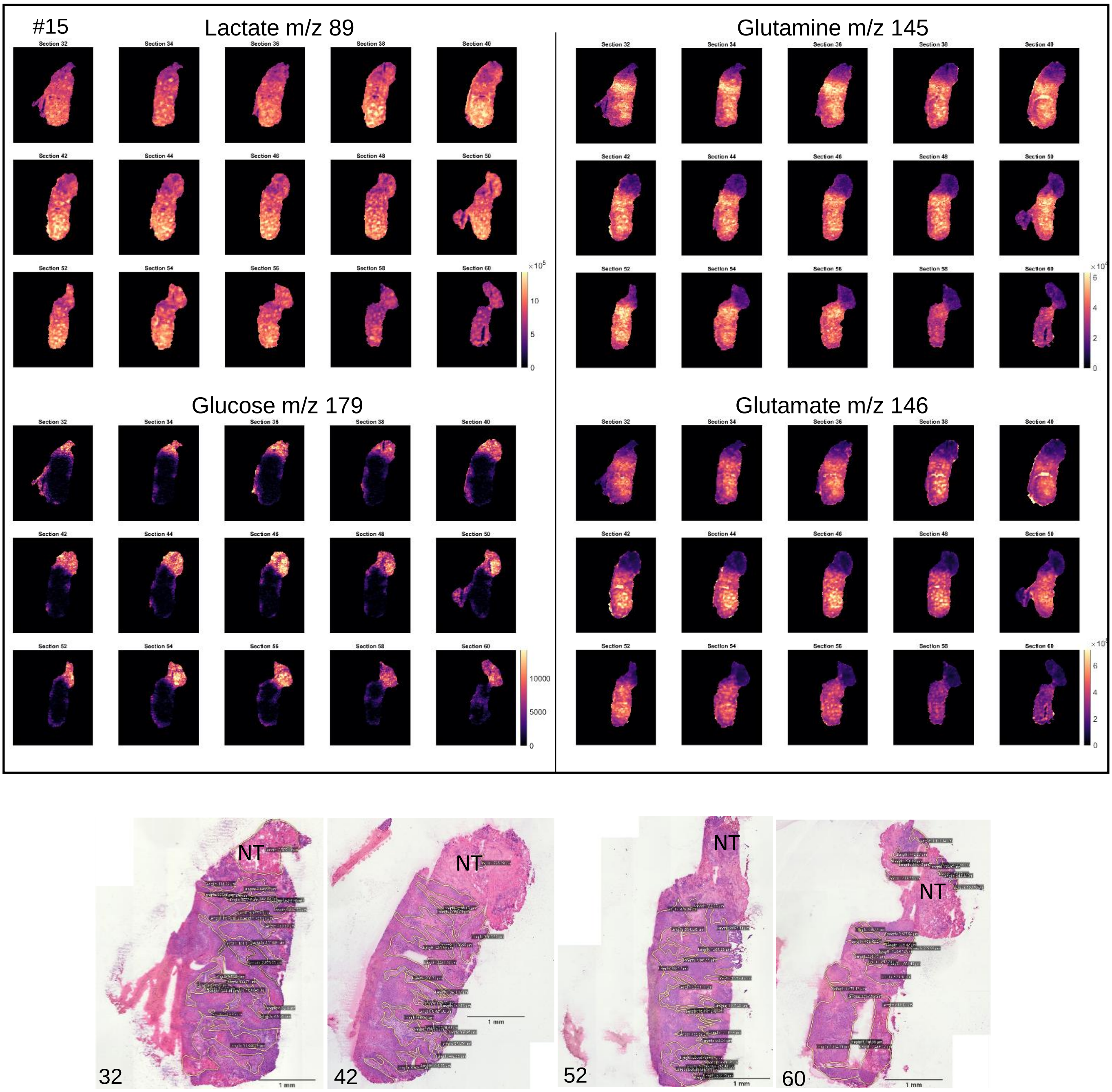


**Supplementary Figure 3. Metabolite distribution in consecutive sections of tumor specimens. (A, B)** Two tumor specimens were flash frozen immediately after harvesting. Selected metabolites were analyzed with spatial resolution of 50 µm by DESI-MRM-MSI in 10 µm thick tumor tissue sections embedded in hydrogel.  For #14 every third section was investigated, for #15 every second section.

HE staining and tissue annotation

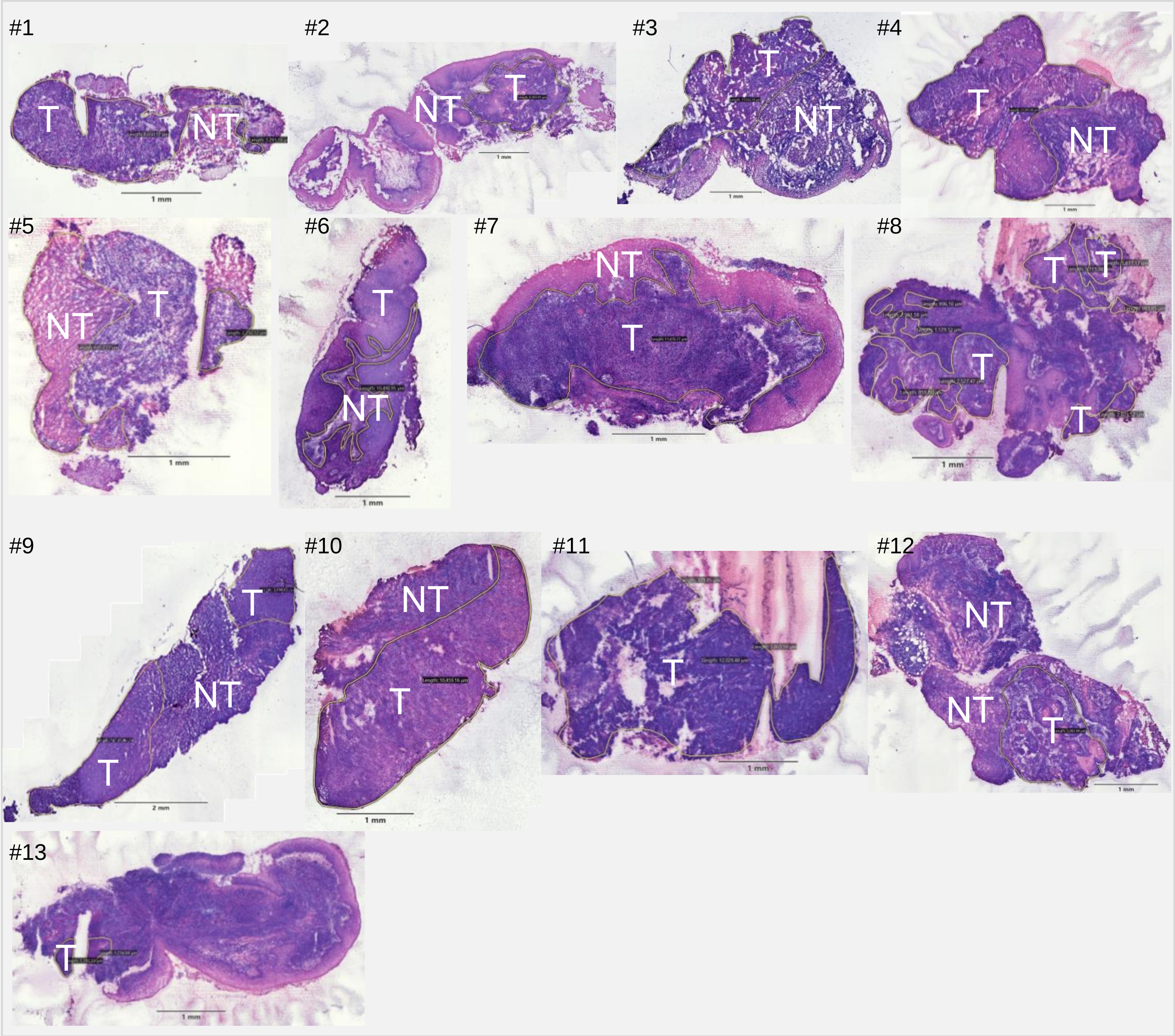


**Supplementary Figure 4. Morphologic characteristics of tumor sections.** H/E stains of the same tumor sections subjected to metabolic imaging and CD3 determination (consecutive section) by IHC. Tumor cell dominated areas (T) are encircled in yellow to separate from non-tumor tissue (NT).

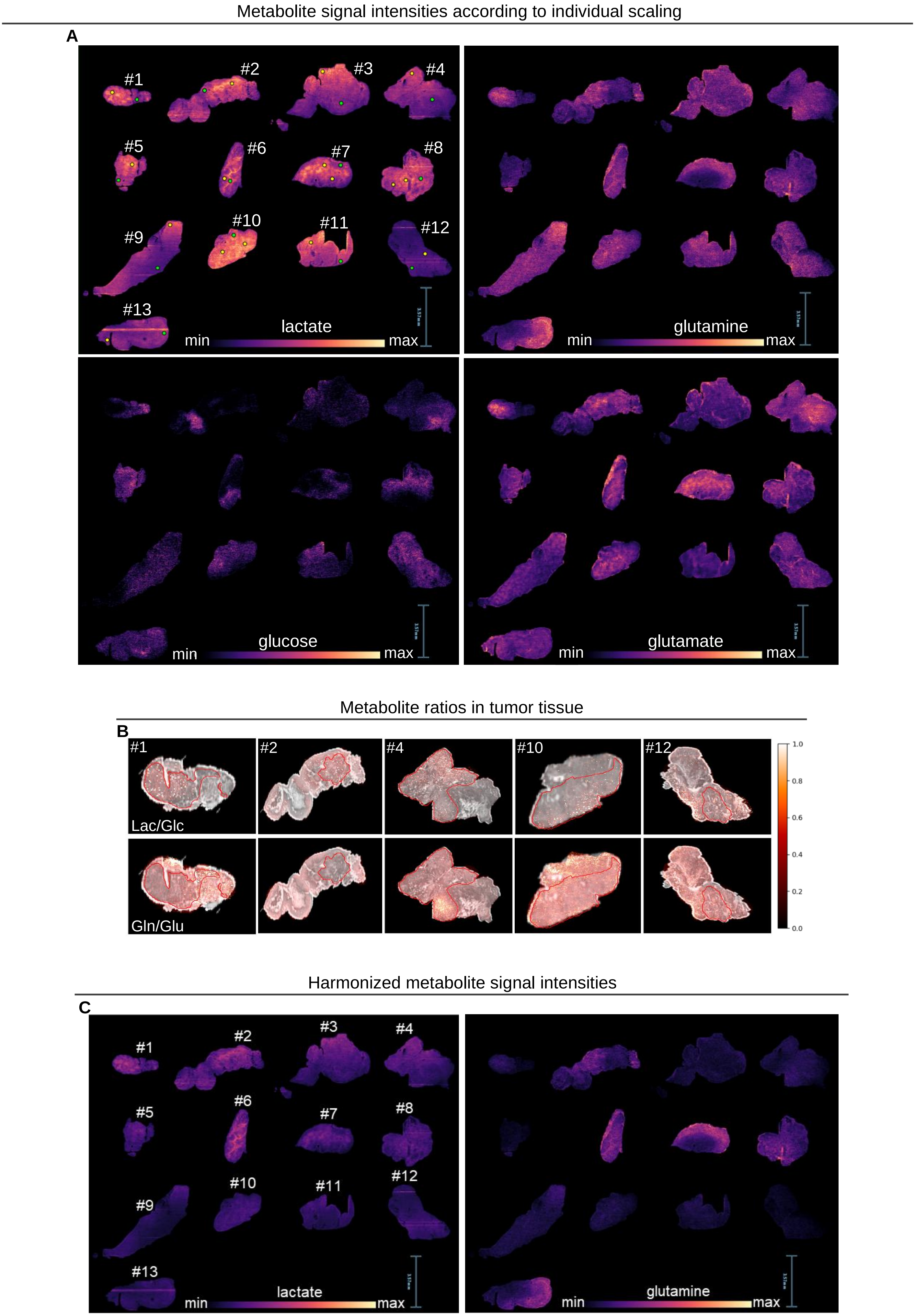
Metabolite signal intensities according to individual scaling
A
#1
#2
#3
#4
#5
#6
#7
#8
#9
#10
#11
#12
#13
lactate
min
max
glutamine
min
max
glucose
min
max
glutamate
min
max
Metabolite ratios in tumor tissue
B
#1
#2
#4
#10
#12
Lac/Glc
Gln/Glu
1.0
0.8
0.6
0.4
0.2
0.0
Harmonized metabolite signal intensities
C
#1
#2
#3
#4
#5
#6
#7
#8
#9
#10
#11
#12
#13
lactate
min
max
glutamine
min
max

**Supplementary Figure 5. Spatial distribution of metabolites.** Selected metabolites were analyzed with spatial resolution of 25 µm by DESI-MRM-MSI in 10 µm thick tumor tissue sections embedded in hydrogel. (**A**) Metabolite intensities are displayed individually scaled for each section to show abundance within a specimen. Regions of interest (ROIs) representative for tumor (T, yellow circles) and non-tumor (NT, green circles) areas within a section for signal intensity extraction are displayed. **(B)** Lactate/glucose and glutamine/glutamate ratios of signal intensities were calculated and co-registered with the morphologic information obtained from H/E stains of the same sections. (**C**) Metabolite intensities are displayed with harmonized scaling across all sections to show abundance between specimens.

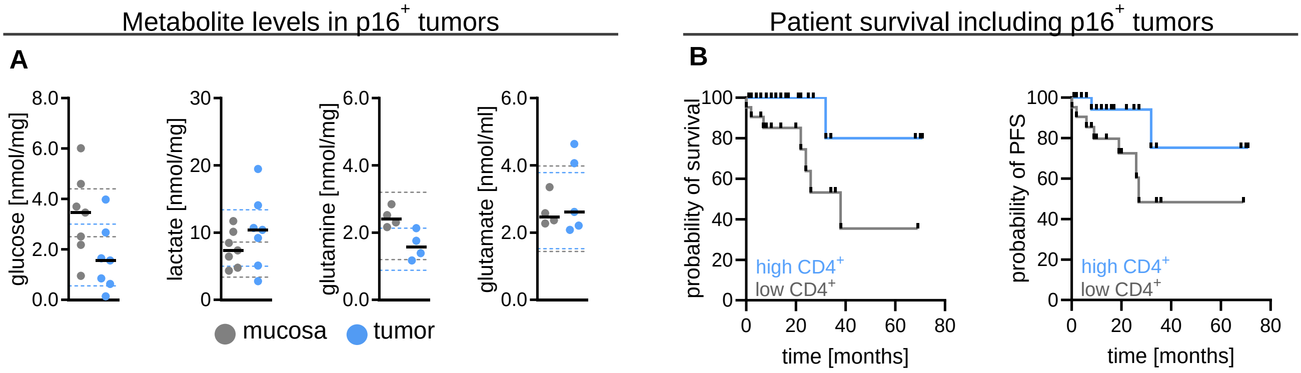


**Supplementary Figure 6. Characteristics of p16⁺ tumors. (A)** Metabolite concentrations in homogenates of p16⁺ tumor tissue and matched mucosa. Tissue was harvested before blood supply cessation and immediately flash frozen in the operating theatre and stored at –80 °C. Shown are single data point and median levels. Dashed lines indicate the 75 % and 25 % percentile of metabolite concentration in p16⁻ tumors (blue) and matched mucosa (grey). **(B)** Clinical data on overall and progression-free survival (PFS) irrespective of p16 status were collected. Shown is OS and PFS dependent on CD4⁺ T cells, high CD4 was defined as percentage above median. Survival was plotted as Kaplan Meier estimation curves.

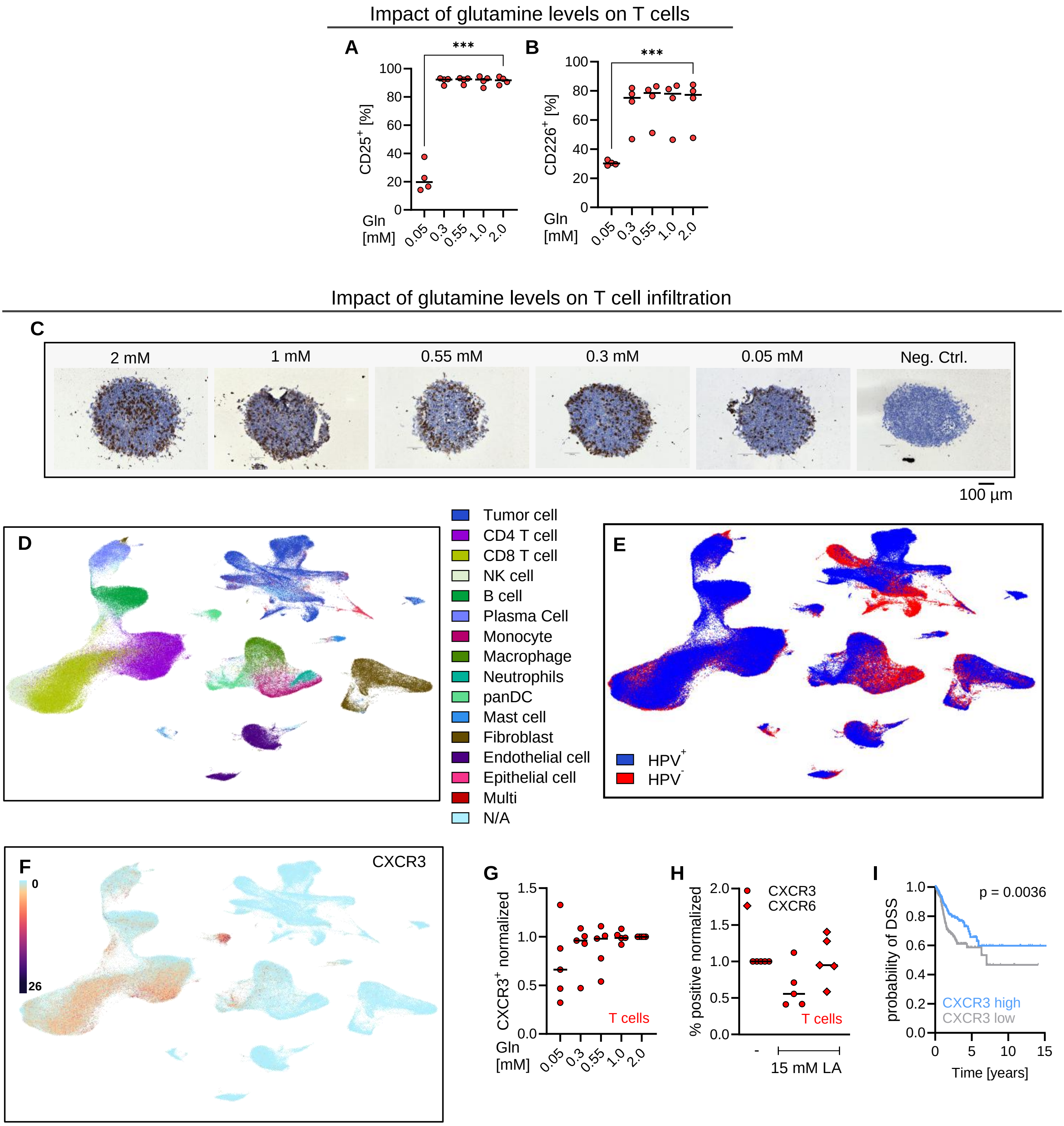


**Supplementary Figure 7. Impact of glutamine restriction on T cell function and infiltration.** (**A, B, G**) T cells were isolated from blood of healthy donors and stimulated with aCD3/CD28 dynabeads in depicted glutamine concentrations. After 48 h of stimulation T cells were stained and the percentage of (**A**) CD25+ and (**B**) CD226+ T cells was determined. (**C**) PCI-15 spheroids were cultured with immune cells in depicted glutamine concentrations. T cells were detected by immunohistochemical analysis of CD3. PCI-15 spheroids without immune cells were used as a negative control for CD3 staining by immunohistochemistry. One representative experiment is shown. **(D – F)** A publicly available HNSCC single-cell RNAseq dataset was analyzed approaching the UCSC cell browser. **(D)** Annotation of cell populations. **(E)** Annotation of HPV status. **(F)** Expression of CXCR3. **(G)** T cells were isolated from blood of healthy donors and stimulated with aCD3/CD28 dynabeads in depicted glutamine concentrations. CXCR3 expression was determined by flow cytometry after 48 h. **(H)** T cells were stimulated with aCD3/CD28 dynabeads in the presence and absence of 15 mM lactic acid. CXCR3 and CXCR6 expression was determined by flow cytometry after 48 h. CXCR6 expression of T cells was determined after 48 h by flow cytometry. **(I)** *CXCR3* expression in relation to disease-specific survival

(DSS) probability was analyzed in a TCGA data set of Head and Neck cancer using the UCSC Xena browser. *CXCR3* high was defined as above median expression. Survival was plotted as Kaplan Meier estimation curves. Significance was calculated applying the log-rank (Mantel-Cox) test, $p<0.05$ was considered as significant. (**A, B, G, H**) Shown are single data points and median levels. Statistical significance was calculated for multiple group comparison using the mixed one-way ANOVA and post-hoc Dunnett's test was used (***$p < 0.001$).

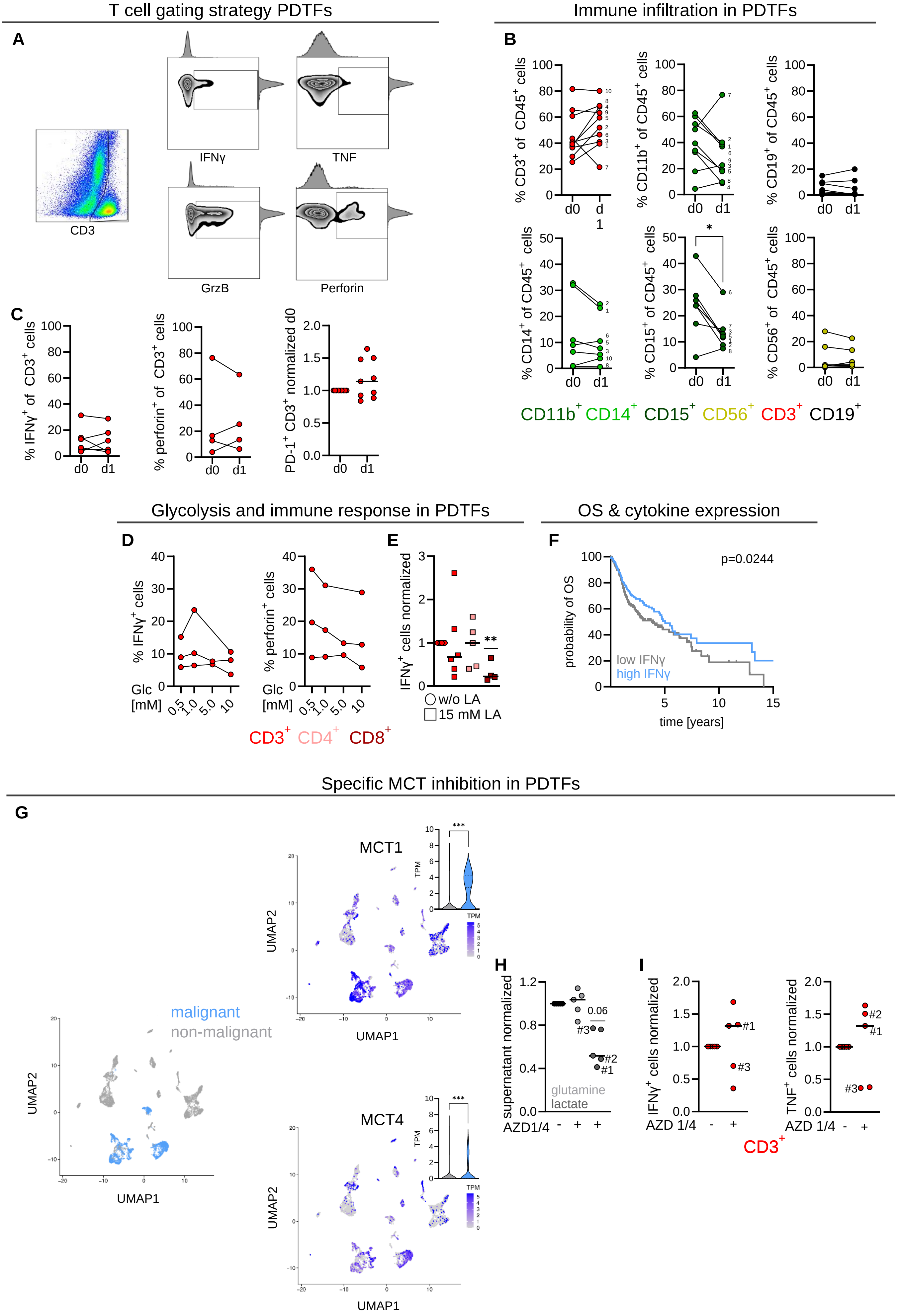

T cell gating strategy PDTFs
Immune infiltration in PDTFs
A
CD3
IFNγ
TNF
GrzB
Perforin
B
% CD3+ of CD45+ cells
% CD11b+ of CD45+ cells
% CD19+ of CD45+ cells
% CD14+ of CD45+ cells
% CD15+ of CD45+ cells
% CD56+ of CD45+ cells
d0
d1
CD11b+ CD14+ CD15+ CD56+ CD3+ CD19+
C
% IFNγ+ of CD3+ cells
% perforin+ of CD3+ cells
PD-1+ CD3+ normalized d0
Glycolysis and immune response in PDTFs
OS & cytokine expression
D
% IFNγ+ cells
% perforin+ cells
Glc [mM]
CD3+ CD4+ CD8+
E
IFNγ+ cells normalized
w/o LA
15 mM LA
F
probability of OS
p=0.0244
low IFNγ
high IFNγ
time [years]
Specific MCT inhibition in PDTFs
G
malignant
non-malignant
UMAP1
UMAP2
MCT1
MCT4
TPM
H
supernatant normalized
glutamine
lactate
AZD1/4
I
IFNγ+ cells normalized
TNF+ cells normalized
AZD 1/4
CD3+

**Supplementary Figure 8. Impact of glycolytic conditions and anti-metabolic targeting on immune cell function.** Tissue biopsies were harvested and immediately transferred to the laboratory. Fragments of approximately 20 mg were prepared, minced, cultivated in culture medium with autologous serum and subjected to indicated conditions. (**A**) Gating strategy for intracellular cytokines is shown. (**B, C**) Biopsies were divided, one part was immediately processed (d0) and the other part was cultured for 24 hours (d1). (**B**) Single cell suspensions were prepared and proportion of immune cell populations was investigated by flow cytometry. Single data points are displayed and statistical significance was calculated using the Wilcoxon test. (**C**) Single cell suspensions were prepared and intracellular cytokine and PD1 expression was investigated by flow cytometry. Single data points are displayed, statistical significance was calculated using the Wilcoxon test or for normalized data the Wilcoxon matched-pairs signed rank test. (**D**) Fragments were cultured in the presence indicated glucose concentrations or **(E)** in the presence or absence of 15 mM lactic acid (LA, 10 mM glucose) for 24 hours, medium was equilibrated in the incubator for at least 3 hours before start of the culture. Single cell suspensions were prepared and intracellular IFNγ and perforin expression determined by flow cytometry. Single data points and median are displayed. Statistics was calculated using for multiple group comparison the Kruskal-Wallis test and post-hoc Dunn's test was used. For normalized data the Wilcoxon matched-pairs signed rank test was used. **(F)** *IFNγ* expression in relation to OS probability was analyzed in a TCGA data set of Head and Neck cancer using the UCSC Xena browser. *IFNγ* high was defined as above median expression. Survival was plotted as Kaplan Meier estimation curves. Significance was calculated applying the log-rank (Mantel-Cox) test, $p<0.05$ was considered as significant. (**G**) UMAP projection of the single cell RNAseq data set of Puram et al. is visualized with malignant and non-malignant cells marked (left). The upper and lower UMAPs show the expression levels of MCT1 and MCT4, respectively, across all cells. The respective violin blots showing raw TPM values are displayed and statistical significance was calculated using the Wilcoxon test,***$p < 0.001$. (**H, I**) Fragments were cultured in the presence or absence of MCT1 and MCT4 inhibitors (1 µM) for 24 hours. (**H**) Glutamine and lactate levels were measured by MS-based methods in supernatants of fragments. (**I**) Single cell suspensions were prepared and intracellular IL-6, IFNγ and TNF determined by flow cytometry in respective immune cell populations. Single data points and median levels are shown and matched specimens are marked by numbers, statistical significance was calculated using the Wilcoxon matched-pairs signed rank test**.**

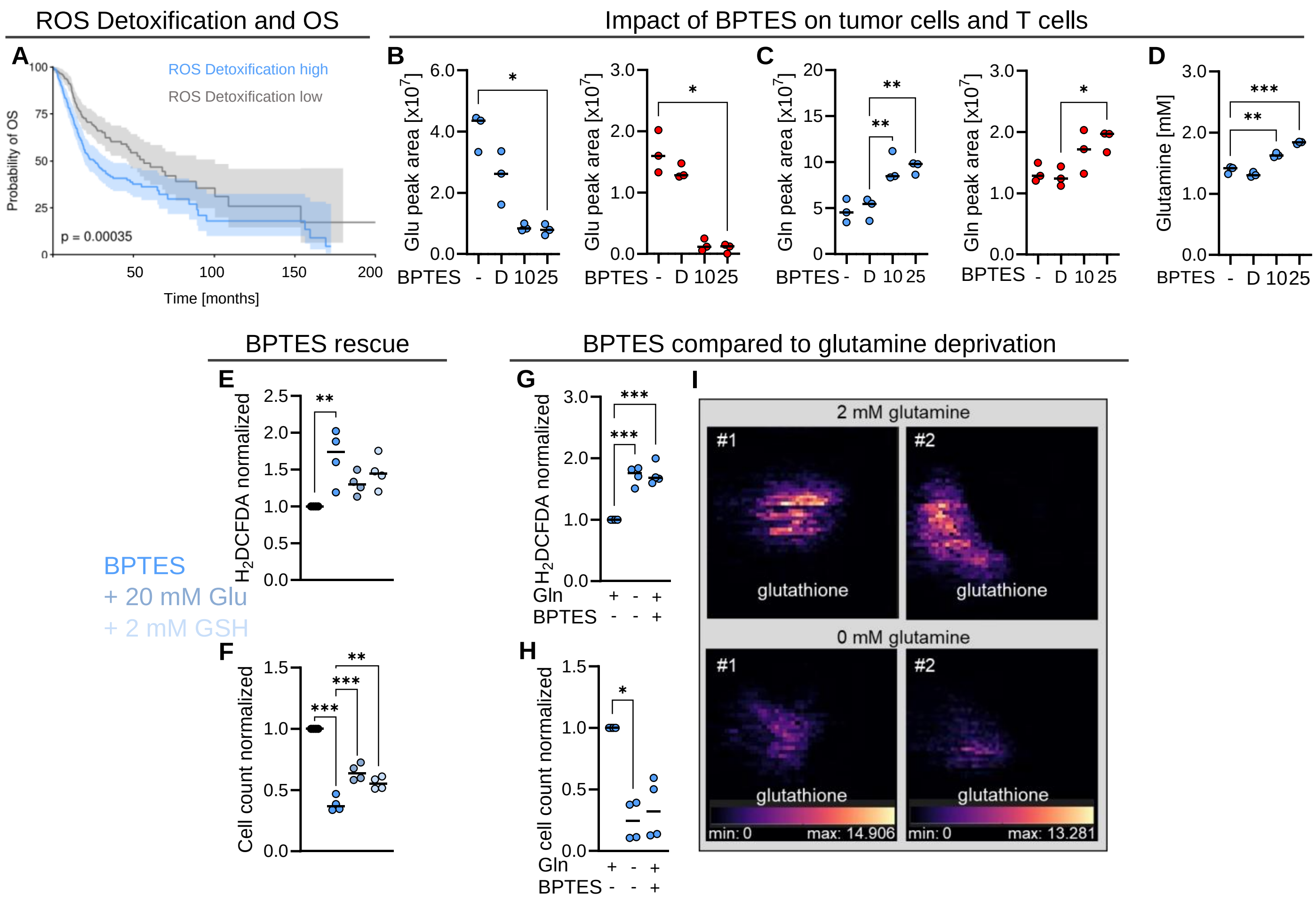


**Supplementary Figure 9. Impact of BPTES on T cells and tumor cells. (A)** Patient survival in relation to expression of genes belonging to the REACTOME_DETOXIFICATION_OF_REACTIVE_OXYGEN_SPECIES (ID R-HSA-3299685) was analyzed. Cutoff was set according to maximally selected rank statistics. **(B-F)** T cells were isolated from blood of healthy donors and stimulated with aCD3/CD28 dynabeads for 48 h in depicted BPTES concentrations [µM]. PCI-15 cells were cultured as described. **(B, C)** Culture medium was supplemented with 2 mM of 13C5 glutamine (PCI-15) or 13C5, 15N2-glutamine (T cells) for 24 h in depicted BPTES concentrations. Intracellular metabolite levels were investigated by mass spectrometry. Integrated peak areas of (**B**) glutamate (Glu) and (**C**) glutamine (Gln) are displayed. (**D**) PCI-15 cells were cultured in the presence of indicated BPTES concentrations for 24 h and glutamine concentration in culture medium was measured by mass spectrometry. (**E, F**) PCI-15 cells were cultivated in the presence and absence of 25 µM BPTES combined with 20 mM Glutamate or 2 mM GSH-MEE for 72 h. **(E)** ROS levels were determined by $H_2$DCFDA staining. **(F)** Cell count was assessed. **(G, H)** PCI-15 cells were cultured in the presence or absence of glutamine or glutamine combined with 10 µM BPTES. After 72 hours **(G)** intracellular ROS levels, determined by staining with $H_2$DCFDA and analyzed by flow cytometry, and **(H)** proliferation, determined with the CASY cell counter, were investigated. (**I**) Glutathione levels in PCI-15 spheroids cultivated with and without glutamine were analyzed with spatial resolution of 25 µm by DESI-MRM-MSI in 10 µm thick sections embedded in hydrogel. (**A-H**) Single data points and median levels are shown, for multiple group comparison the Friedman and post-hoc Dunnett's test or the Kruskal Wallis test and post-hoc Dunn's was used (*p < 0.05, **p < 0.01, ***p < 0.001)**.**

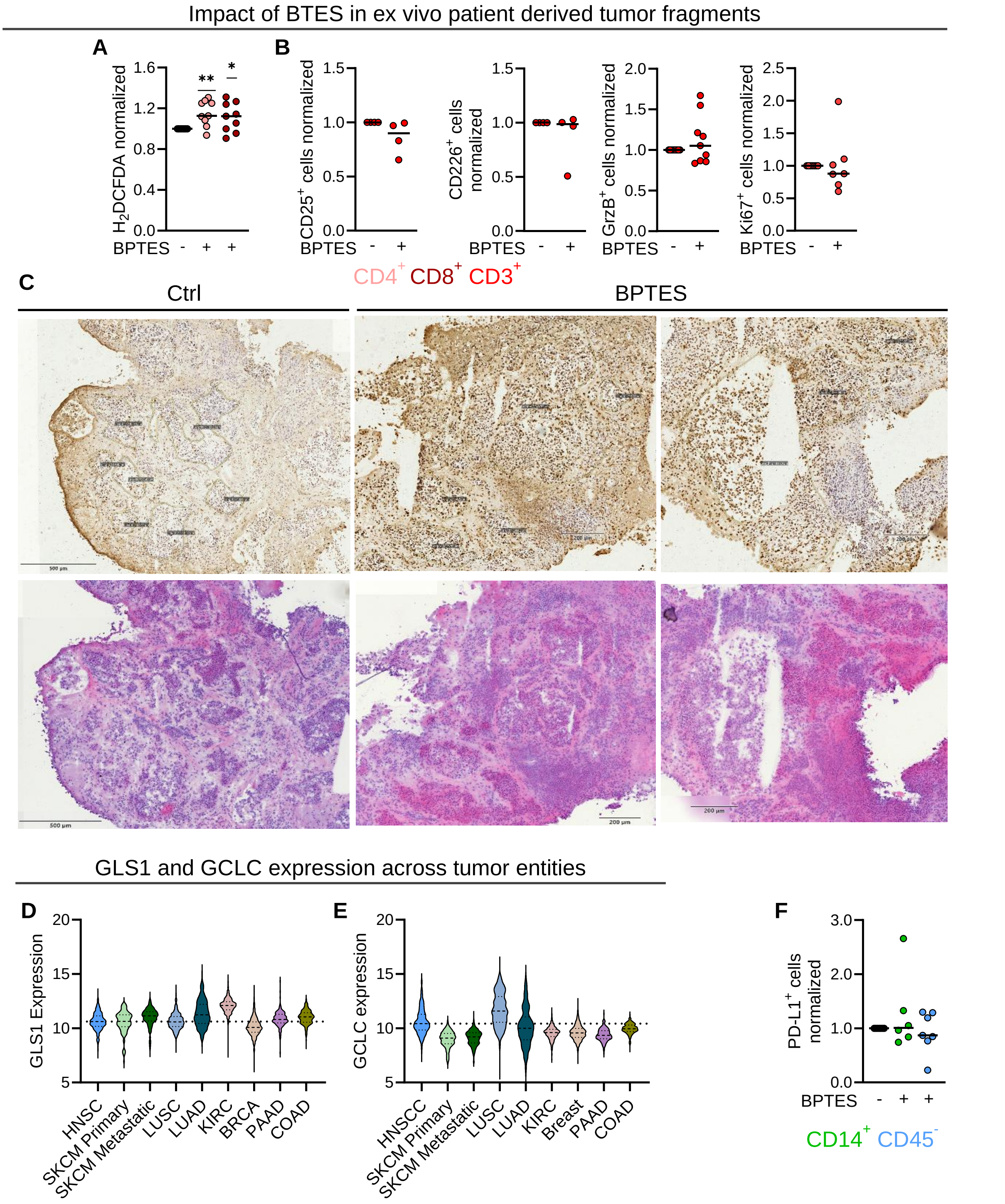


**Supplementary Figure 10. Impact of BPTES on patient derived tumor fragments.** Tissue biopsies were harvested and immediately transferred to the laboratory. Fragments of approximately 20 mg were prepared, minced, cultivated in culture medium with autologous serum and subjected to 10 µM of BPTES. Single cell suspensions were prepared and stained for immune cell population specific surface markers. (**A**) Intracellular ROS levels were determined in $CD4^+$ and $CD8^+$ T cell populations by staining with $H_2$DCFDA and analyzed by flow cytometry. (**B**) T cells were stained and the percentage of CD25+, $CD226^+$, Granzyme $B^+$ ($GrzB^+$) and $Ki67^+$ T cells was determined by flow cytometry. (**C**) Fragments were embedded in hydrogel, 10 µm sections prepared and stained for apoptosis using the TUNEL assay. Apoptotic nuclei are stained brown, viable nuclei are of blue colour. Tumor-cell dominated areas are encircles. **(D)** GLS1 and **(E)** GCLC expression across tumor entities was analyzed in the TCGA Pan-Cancer dataset. **(F)** The percentage of PD-$L1^+$ $CD14^+$ and $CD45^-$ cells in BPTES-treated PDTFs was determined by flow cytometry. (**A, B, F**) Single data points and median levels are shown, for normalized data comparison the Wilcoxon matched-pairs signed rank test was used (*$p < 0.05$, **$p < 0.01$)**.**

**Supplementary Table 1. Patient cohort characteristics.**

| patient characteristics | No. (n=244) | [%] |
|---|---|---|
| **Age at surgery (median)** | 64 | |
| **Sex** | | |
| Male | 194 | 79.5 |
| Female | 49 | 20.1 |
| **Tumor localsation** | | |
| Oral cavity | 61 | 25.0 |
| Oropharynx | 94 | 38.5 |
| Larynx/Hypopharynx | 79 | 32.4 |
| Multiple localisation | 10 | 4.1 |
| **HPV status** | | |
| p16 positive | 33 | 13.5 |
| **T category** | | |
| T1/2 | 114 | 46.7 |
| T3/4 | 129 | 52.9 |
| **N category** | | |
| N0 | 110 | 45.1 |
| N1 | 41 | 16.8 |
| N2 | 36 | 14.8 |
| N3 | 6 | 2.5 |
| unknown | 27 | 11.1 |
| **UICC stage (TNM eighth edition)** | | |
| I | 33 | 13.5 |
| II | 53 | 21.7 |
| III | 46 | 18.9 |
| IVA | 66 | 27.0 |
| IVB | 34 | 13.9 |
| IVC | 8 | 3.3 |
| unknown | 3 | 1.2 |
| **Tobacco smoking status** | | |
| yes | 182 | 74.6 |
| no | 55 | 22.5 |
| unknown | 6 | 2.5 |
| **Alcohol consumption status** | | |
| yes | 123 | 50.4 |
| no | 114 | 46.7 |
| unknown | 6 | 2.5 |
| **BMI** | | |
| overweight (> 24.9) | 112 | 45.9 |
| normal weight (18.5-24.9) | 109 | 44.7 |
| underweight (< 18.5) | 19 | 7.8 |
| unknown | 3 | 1.2 |
| **Status post local radiatio** | | |
| yes | 45 | 18.4 |
| no | 195 | 79.9 |
| unknown | 2 | 0.8 |
| **Deceased patients** | 43 | 17.6 |

**Supplementary Table 2. Concentrations of carbon metabolites determined in tissue and ISF of tumor and matched mucosa.**

| | **tumor (tissue)** | | **mucosa (tissue)** | | | | **tumor (ISF)** | | **mucosa (ISF)** | | | |
|---|---|---|---|---|---|---|---|---|---|---|---|---|
| **metabolite [mM]** | **median** | **SD** | **median** | **SD** | **p-value** | **adjusted p-value** | **median** | **SD** | **median** | **SD** | **p-value** | **adjusted p-value** |
| lactate | 7.11 | 5.02 | 5.23 | 2.96 | **0.0043** | **0.0067** | 10.98 | 5.76 | 5.55 | 4.18 | **0.0021** | **0.0155** |
| glucose | 1.86 | 1.43 | 3.38 | 1.58 | **0.0001** | **0.0017** | 1.50 | 1.74 | 4.32 | 2.58 | **0.00001** | **0.0017** |
| glutamine | 1.43 | 0.95 | 2.32 | 1.23 | **0.0037** | **0.0050** | 1.24 | 0.67 | 1.53 | 0.81 | 0.0709 | n.s. |
| glutamate | 2.58 | 1.34 | 2.14 | 1.61 | 0.74 | n.s. | 2.34 | 0.92 | 1.13 | 1.10 | **0.0010** | **0.0052** |
| citrate | 0.22 | 0.21 | 0.28 | 2.40 | **0.0104** | **0.0067** | 0.09 | 1.50 | 0.08 | 0.04 | 0.56 | n.s. |
| 2-hydroxyglutarate | 0.02 | 0.02 | 0.01 | 0.03 | 0.19 | n.s. | n.d. | n.d. | n.d. | n.d. | n.d. | n.s. |
| α-ketoglutarate | 0.04 | 0.03 | 0.04 | 0.04 | 0.57 | n.s. | 0.11 | 0.43 | 0.06 | 0.48 | 0.64 | n.s. |
| pyruvate | 0.08 | 0.23 | 0.09 | 0.41 | 0.59 | n.s. | 0.30 | 0.32 | 0.20 | 0.11 | 0.08 | n.s. |
| succinate | 0.18 | 0.58 | 0.13 | 0.20 | 0.29 | n.s. | 0.19 | 0.94 | 0.08 | 0.08 | **0.0186** | **0.024** |
| malate | 0.29 | 0.24 | 0.35 | 6.38 | 0.14 | n.s. | 0.34 | 0.27 | 0.25 | 0.20 | 0.11 | n.s. |
| fumarate | 0.10 | 0.07 | 0.10 | 0.06 | 0.94 | n.s. | 0.09 | 0.24 | 0.06 | 0.34 | 0.99 | n.s. |
| threonine | 0.36 | 0.21 | 0.31 | 0.15 | 0.22 | n.s. | 0.36 | 0.20 | 0.24 | 0.11 | **0.0012** | **0.0069** |
| alanine | 1.11 | 0.93 | 1.14 | 0.86 | 0.97 | n.s. | 1.39 | 0.82 | 0.86 | 0.48 | **0.05** | n.s. |
| proline | 0.51 | 0.45 | 0.26 | 0.39 | **0.0002** | **0.0033** | 0.57 | 0.39 | 0.31 | 0.13 | **0.0023** | **0.0172** |
| methionine | 0.05 | 0.03 | 0.05 | 0.15 | 0.0947 | n.s. | 0.08 | 0.04 | 0.04 | 0.02 | **0.0013** | **0.009** |
| aspartate | 0.60 | 0.39 | 0.53 | 0.61 | 0.18 | n.s. | 0.69 | 0.45 | 0.23 | 0.43 | **0.0038** | **0.0190** |
| valine | 0.36 | 0.21 | 0.34 | 0.15 | 0.29 | n.s. | 0.45 | 0.17 | 0.32 | 0.14 | **0.0254** | n.s. |
| histidine | 0.17 | 0.09 | 0.16 | 0.07 | 0.79 | n.s. | 0.19 | 0.10 | 0.14 | 0.07 | 0.21 | n.s. |
| lysine | 0.32 | 0.23 | 0.36 | 0.17 | 0.22 | n.s. | 0.42 | 0.27 | 0.33 | 0.16 | **0.03** | n.s. |
| leucine | 0.29 | 0.11 | 0.26 | 0.10 | 0.10 | n.s. | 0.33 | 0.15 | 0.22 | 0.10 | **0.0020** | **0.0138** |
| asparagine | 0.23 | 0.10 | 0.12 | 0.06 | **0.0109** | **0.0083** | 0.16 | 0.07 | 0.08 | 0.04 | **0.0013** | **0.0103** |
| glycine | 1.64 | 0.90 | 1.59 | 1.36 | 0.76 | n.s. | 1.57 | 1.03 | 0.83 | 0.72 | **0.0047** | **0.0207** |
| arginine | 0.22 | 0.11 | 0.24 | 0.13 | 0.10 | n.s. | 0.21 | 0.12 | 0.17 | 0.07 | 0.08 | n.s. |
| serine | 0.40 | 0.21 | 0.38 | 0.23 | 0.53 | n.s. | 0.40 | 0.21 | 0.30 | 0.22 | 0.06 | n.s. |
| ornithine | 0.08 | 0.04 | 0.09 | 0.07 | 0.12 | n.s. | 0.12 | 0.08 | 0.14 | 0.04 | 0.39 | n.s. |
| phenylalanine | 0.15 | 0.10 | 0.14 | 0.05 | 0.26 | n.s. | 0.20 | 0.09 | 0.10 | 0.05 | **0.0005** | **0.003** |
| isoleucine | 0.13 | 0.08 | 0.10 | 0.05 | **0.0198** | n.s. | 0.16 | 0.07 | 0.10 | 0.04 | **0.0014** | **0.0121** |
| tyrosine | 0.13 | 0.12 | 0.19 | 0.06 | 0.64 | n.s. | 0.19 | 0.08 | 0.11 | 0.04 | **0.0062** | **0.022** |
| tryptophan | 0.04 | 0.03 | 0.04 | 0.02 | 0.54 | n.s. | 0.04 | 0.02 | 0.04 | 0.02 | 0.80 | n.s. |
| 3-hydroxybutyrate | 0.58 | 0.55 | 0.53 | 0.49 | 0.55 | n.s. | 0.61 | 0.60 | 0.57 | 0.71 | 0.59 | n.s. |

Metabolite concentrations were determined in tissue homogenates and ISF of matched tumor and mucosa samples. Metabolites in tissue was determined in nmol/mg, which corresponds to mM. Displayed is the median and the p-value. For tissue samples, 37 (organic acids) and 30 (amino acids) matched specimens were analysed except for 2-hydroxyglutarate (n = 12), α-ketoglutarate (n = 24), asparagine (n = 15) and tyrosine (n = 13), in the other samples the respective metabolites were below limit of quantification. For ISF, 23 matched samples were analysed except for α-ketoglutarate, citrate, asparagine (each n = 12) as well as succinate and pyruvate (each n = 15) and fumarate (n = 10). Normality was tested applying the D'Agostino & Pearson and Shapiro-Wilk test. Statistical significance was calculated using a paired t-test or the Wilcoxon matched-pairs signed rank test (exact p-values given). To adjust for multiple testing, the Benjamini-Hochberg procedure was applied to control the FDR at 0.05 (adjusted p-values).

**Supplementary Table 3. Signal intensities for respective metabolites from regions of interest.**

| sample ID | ROI sum peak | lactate | glucose | glutamine | glutamate |
|---|---|---|---|---|---|
| #1_T | 54316908 | 25688195 | 3786 | 1092121 | 27532902 |
| #1_NT | 17461240 | 13037210 | 98918 | 322664 | 4002576 |
| #2_T | 30650353 | 24021354 | 3277 | 889337 | 5736497 |
| #2_NT | 20041397 | 13442001 | 1966 | 1599949 | 4997581 |
| #3_T | 31378872 | 22361662 | 36562 | 600889 | 8379882 |
| #3_NT | 14704660 | 9445822 | 14423 | 487845 | 4756681 |
| #4_T | 27377238 | 19505381 | 59702 | 761195 | 7051085 |
| #4_NT | 19011327 | 9609541 | 127458 | 803166 | 8471295 |
| #5_T | 27384548 | 18700457 | 167669 | 347054 | 8169505 |
| #5_NT | 14571478 | 10284196 | 92224 | 344201 | 3850994 |
| #6_T | 30248006 | 17179400 | 18093 | 1627709 | 11422926 |
| #6_NT_fibro. | 45305346 | 26826341 | 224561 | 2323843 | 15930741 |
| #7_T_high_Lac | 35863099 | 16633729 | 60887 | 2255633 | 16912989 |
| #7_T_low_Lac | 23094044 | 10800615 | 21428 | 930521 | 11341605 |
| #7_NT | 21510258 | 9085281 | 6200 | 2499570 | 9919323 |
| #8_T_high_Lac | 27905746 | 14932694 | 37496 | 1769841 | 11165847 |
| #8_T_low_Lac | 22428643 | 10503748 | 30875 | 1555172 | 10338968 |
| #8_NT | 20643277 | 12216529 | 90604 | 1040876 | 7295398 |
| #9_T | 28515903 | 13868722 | 30551 | 1072468 | 13544289 |
| #9_NT | 13921712 | 5206567 | 47681 | 727912 | 7939693 |
| #10_T_high_Gln | 29677666 | 12411318 | 143258 | 986995 | 16136222 |
| #10_T_low_Gln | 22630805 | 11840175 | 45253 | 598913 | 10146596 |
| #10_NT | 14590029 | 8484132 | 69211 | 522342 | 5514475 |
| #11_T_high_Gln | 25084770 | 11374466 | 102574 | 793062 | 12814800 |
| #11_T_low_Gln | 19525163 | 9195475 | 40802 | 450904 | 9838115 |
| #12_T | 15855822 | 10189278 | 161930 | 247163 | 5257584 |
| #12_NT | 18221034 | 12009334 | 66848 | 452211 | 5692778 |
| #13_T | 18503235 | 9055321 | 49535 | 683948 | 8714562 |
| #13_NT | 23632388 | 11616179 | 87732 | 2472907 | 9455706 |

Regions of interest (ROIs, 70 scans, each scan 25 µm to 25 µm) representative of tumor (T, yellow circle) and non-tumor tissue (NT, green circle) were defined (Supplementary Figure 2D), scaling for each metabolite was harmonized between samples and mean signal intensities retracted from the ROIs. Abbreviations used: lactate (Lac), glutamine (Gln), tumor (T), non-tumor (NT).

**Supplementary Material and Methods**

*Details on study participants and patient enrollment*

Patients underwent diagnostic panendoscopy of the upper air and feed paths, primary or salvage surgical resection of the tumor including a neck dissection ipsilateral or on both neck sides or, instead of surgery, received a primary radio-(chemo) therapy based on clinical, ultrasound and further radiologic findings as well as histological confirmation by means of biopsy. Stage classification was performed according to the Union international contre le cancer (UICC) guidelines in the 8$^{th}$ edition (1). The treatment of HNSCC patients was determined by the interdisciplinary tumor board for head and neck malignancies, University Hospital Regensburg, Germany, and not influenced by study inclusion. Disease relapse was defined as tumor or metastatic recurrence by radiologic evidence with clinical correlation and histologic confirmation by biopsy. Patients' death was documented from medical records and the Clinical Cancer Registry of the Tumor Center-Institute for Quality Management and Health Services Research, University of Regensburg, Germany (cut-off date was set at 01.01.2024).

*Immune cell isolation from blood and tissue samples*

Blood leucocytes were isolated from lithium heparin blood by ACK-lysis. 20 mL ACK buffer 1x (6x: 0.155 M $NH_4Cl$, 0.01 M $KHCO_3$ and 0.1 mM EDTA) were added to 1 ml blood and incubated for 5 min at room temperature. 30 mL PBS supplemented with 2 % fetal calf serum (FCS, heat-inactivated, Millipore Sigma; FACS buffer) were added and cells were centrifuged at 515 g for 5 min at 4 °C. The supernatant was discarded and the ACK-lysis was repeated using 10 mL ACK buffer and 20 mL FACS buffer per sample. If erythrocytes were still visible in the pellet, a third lysis step (5 mL ACK and 10 mL FACS buffer) was performed. Finally, cells were pooled and washed in 10 mL FACS buffer, resuspended in FACS buffer and cell counting was performed using the Casy system (Casy® Modell TT, Omni Life Science).

Tumor samples and corresponding mucosa were transferred into RPMI1640 medium (Thermo Fisher Scientific), supplemented with 10 % FCS, 2 mM L-glutamine, penicillin (50 U/mL, Thermo Fisher Scientific) and streptomycin (50 µg/mL, Thermo Fisher Scientific), collagenase type 4 (4160 IU; Worthington) and 10 mg/mL DNAse I (Sigma) and further minced with a sterile scalpel. The mixture was incubated at 37 °C and 5 % $CO_2$ for 2 h for digestion. Following, the tissue sample was pushed through a 70 µm nylon mesh (Falcon) and washed twice with medium. If required, an ACK-lysis was performed to remove residual erythrocytes (3 mL ACK and 10 mL FACS buffer). Cells were washed, resuspended in FACS buffer, counted, and further processed for flow cytometry.

*Metabolite profiling*

*Determination of central carbon metabolites and amino acids in tissue homogenates and interstitial fluid*

Extraction of metabolites from tumor tissues and ISF was performed as recently described (2). An aliquot of the resulting aqueous extract was used for analysis of intermediates of glycolysis and TCA cycle by gas chromatography-mass spectrometry after derivatization as described (2). A 10 µL aliquot of the final extract was utilized for amino acid analysis by high performance liquid chromatography – electrospray ionization – tandem mass spectrometry (ExionLC AD with Triple Quad 6500+, AB SCIEX). For analysis, amino acids were subjected to derivatization with propyl chloroformate/propanol (3) using 17.4 % propyl chloroformate and 82.6 % isooctane as reagent 2 in contrast to original protocol.

*Spatial metabolic imaging of tissue and tumor spheroid sections*

The bottom of embedding molds (bottom dimensions 8 x 8 mm; Kisker Biotech) was covered with hydrogel matrix and allowed to slightly freeze. The frozen tumor tissue was placed at the center of the mold and completely covered with hydrogel matrix. Embedded tissue was stored at -80 °C until sectioning. Sectioning of the samples was performed with the cryostat kept at -35 °C using SuperfrostTM Plus Adhesion Microscope slides (Epredia). Slides were washed with distilled water and isopropanol before usage. Sections of 10 µm thickness were prepared, stored at -80°C and analyzed within 48 h. To optimize the parameters for DESI-MRM-MSI, 10 µM solutions of the metabolites in question were directly injected via the ESI source into a TQ-XS triple quadrupole mass spectrometer (Waters) to tune Cone Voltage (CV) and Collision Energy (CE) in negative ion mode. Optimized transition parameters are given below.

| Metabolites | Precursor ion (m/z) | Product ion (m/z) | CV (V) | CE (eV) |
|---|---|---|---|---|
| Lactate | 89 | 43 | 5.0 | 10.0 |
| Glutamine | 145 | 109 | 28.0 | 16.0 |
| Glutamate | 146 | 102 | 20.0 | 20.0 |
| Glucose | 179 | 89 | 28.0 | 10.0 |
| Glutathione | 306 | 143 | 20.0 | 25.0 |

The setup for DESI-MRM-MSI consisted of a TQ-XS mass spectrometer coupled to a Prosolia DESI stage with a high-performance sprayer (Waters, Wilmslow, UK). Sprayer solvent flow was provided by a nanoACQUITY Binary Solvent Manger (Waters). A 95:5 (v/v) mixture of LC-MS grade methanol (Sigma-Aldrich) and water (Purelab® flex) was used as DESI solvent at a flow rate of 2 µL/min with a nitrogen nebulizing gas pressure of 1 bar. Sprayer angle was positioned at 75°. DESI inlet capillary heating was set to 350 °C and sprayer capillary voltage was set to 550 V. The HNSCC samples were vacuum dried for 5 min before measurement. DESI-imaging for 1 MRM transition per metabolite was performed at 25 µm spatial resolution for Figure 3 and 8 and with 50 µm resolution for Figure 4 in in a scan time of 10 Hz and negative ion mode. Images were acquired with HDI 1.7 and MassLynx 4.1 (Waters). Image processing was also done with HDI 1.7.

For imaging of spheroid sections, the same setup as for tissue sections was used.

*3D sample preprocessing and coregistration*

The initial coregistration was between each MS section's lactate ion image and its corresponding H&E-stained image. Both images were scaled between 0-1 and an initial threshold of 0.7 was used to differentiate tissue from background; these thresholds were manually altered if required. An affine transformation was determined that maximised the overlap between binary masks, from which the warped H&E image was generated.

For coregistration of serial sections, a rigid transformation of the n+1th binarized lactate image was determined with reference to that from the nth section. To generate the 3D stack, images were then sequentially warped from the second section onwards by multiplying a section's

transform by all preceding transformations. The same rigid transformation was applied to each of the MS images. For the optical image stack (with higher resolution images), the procedure is the same, but the transformations are scaled according to the image size ratio between the MS and H&E images.
To extract the annotations from an H&E image, it was first converted to the L*a*b* colour space. Values of b* greater than 25 were identified as annotations and the image was skeletonised. Any individual annotations consisting of fewer than 20 pixels were discarded. The remaining pixel coordinates were rescaled to match the MS image resolution.
For image visualisation purposes, the aura surrounding each tissue section was determined and removed. The lactate ion was used to identify the approximate tissue outline. The aura was identified as those pixels with a total intensity greater than the 98th (sample 216) or 95th (sample 239) percentile surrounding the lactate-determine tissue mask. The mask was morphologically closed (filter radius = 2) and then eroded (filter radius = 1) to further minimise the appearance of erroneous high intensity pixels.
For image visualisation purposes, the maximum intensity for each ion is constant across all sections (in image arrays and animations). That value is the maximum value determined from the 99.5th percentile intensity from each section.

*Hematoxylin/Eosin (H/E) staining*
After DESI-MRM-MSI, slices were subjected to H/E staining. After thawing and drying, slides were fixed with 4 % paraformaldehyde for 10 min at 4 °C. After two washes with water for 5 min each, slices were dipped in hematoxylin solution. Excess hematoxylin was removed by washing with distilled water. Subsequently, slides were rinsed in running tap water for 10 min for staining intensification. Following, slices were incubated in eosin for 3-6 s. Slides were dehydrated (96 % ethanol, 2x 100 % ethanol, 2x xylol, 8 min each), and mounted with glass coverslips in permanent mounting media Entellan® Neu (VWR). Images were recorded using the M8 Microscope and Scanner (Precipoint). Images of all tumor sections included in the paper are shown in Supplementary Figure 6.

*Immunohistochemistry*
IHC was performed on cryosections of tumor tissue, embedded spheroids and co-cultures as prepared for DESI-MRM-MSI. Prior to immunohistochemistry staining, tissue sections mounted on glass slides were thawed, dried and fixed in 4 % paraformaldehyde for 10 min at 4 °C. For antigen-retrieval, slices were heated to 95 °C for 30 min in Tris-EDTA buffer (Biozol,). After cooling down for 30 min, sections were washed with distilled water. Endogenous peroxidase activity was blocked by incubation in Dako REAL Peroxidase Blocking solution for 5 min. Sections were washed with Dako wash buffer and subsequently stained for CD3 (1:50, 30 min). After renewed washing, labelled Polymer (Histofine ® simple stain AP anti-rabbit, Nichirei Biosciences) was added for 30 min, followed by a 5-min washing step and chromogenic development applying DAB (Liquid DAB+ 2-component system, Dako) according to manufacturer's instructions (reaction time 10 min). After slide washing with water and counterstaining with hemalaun, slides were rinsed with tap water, dehydrated (96 % ethanol, 2x 100 % ethanol, 2x xylol, 8 min each), and mounted with glass coverslips in permanent mounting media Entellan® Neu. Images were recorded using the M8 Microscope and Scanner.

*TUNEL assay*

To assess apoptosis in tumor tissue sections, TUNEL staining was performed using the TUNEL In Situ Apoptosis Kit according to the manufacturer's instructions. The reaction was visualized using DAB as chromogen, resulting in brown staining of apoptotic nuclei. Sections were subsequently counterstained with Mayer's hematoxylin to visualize viable cell nuclei in purple/blue. Images were acquired using M8 Microscope and Scanner. The apoptosis rate was calculated as the percentage of DAB-positive (TUNEL$^+$) nuclei relative to the total number of hematoxylin-stained nuclei. Image analysis was performed using ImageJ (version 1.54p, National Institutes of Health, USA).

*Primer sequences for Quantitative Real-Time PCR*

| Gene | Forward sequence 5' → 3' | Reverse sequence 5' → 3' |
|---|---|---|
| *18S rRNA* | ACCGATTGGATGGTTTAGTGAG | CCTACGGAAACCTTGTTACGAC |
| *GLUT1* | AACTCTTCAGCCAGGGTCCAC | CACAGTGAAGATGATGAAGACGTAGGG |
| *LDHA* | GGTTGGTGCTGTTGGCATGG | TGCCCCAGCCGTGATAATGA |
| *MCT1* | CAGCTTCTTTCTGTAACACCGT | GTCGCCTCTTGTAGAAATACTTGCC |
| *MCT4* | TGCCCCAGCCGTGATAATGA | CATCCAGGAGTTTGCCTCCCGA |
| *GLS* | GCAGTCTGGAGGAAAGGTTGCA | TACACAGGACTGAAGACAGAAGGGA |
| *SLC7A5* | GCCCTCCATCCTCTCCATGATCCA | TGAAGAAGCTGAAGAAGTTGATGACGGA |
| *SLC7A8* | TGGAGGCTGGAACTTTCTGAATTACGTG | AGATGAAGATGGCTCTGGGAAGGT |

*Analysis of metabolic dependencies of tumor cell lines*

For analysis of glutamine dependency, cells were subjected to L-glutamine concentrations or 5 nM oligomycin (Sigma-Aldrich). For analysis of BPTES effects, medium was replaced by T cell medium (containing 10 % AB serum) with and without 10 µM and 25 µM BPTES and respective DMSO control. For supplementation experiments, cells were additionally treated with 20 mM glutamate or 2 mM glutathione-monoethylester (GSH-MEE; both Sigma-Aldrich). To ensure stability of GSH-MEE, treatment stocks were freshly prepared directly before addition. For analysis of MHCI expression, cells were treated with different concentrations of human recombinant IFNγ (Peprotech, Thermo Fisher Scientific).

After 72 h (metabolic dependencies) and 24 h (MHCI expression) incubation, cells were washed with PBS, detached with 500 µL Trypsin/EDTA (Thermo Fisher Scientific), and subjected to further analyses.

*Generation of GLS1$^{-/-}$ PCI-15 cells*

PCI15 GLS1-/- cells were generated using CRISPR/Cas9. Three publicly available crRNA sequences directed against GLS1 were selected to limit off-target effects (crRNA1 GATTGCGAACGTCTGATCCC; crRNA2 CGCCGGCACAGGGTGACCAA; crRNA3 AGCTCCTGCAAGATCTCCGA; ordered at Integrated DNA Technologies). As a control, three scrambled crRNAs (at Integrated DNA Technologies) were used. To prepare aligned gRNA complexes, crRNA and tracrRNA were mixed in IDTE buffer (final concentration 50 µM each) and incubated for 5 minutes at 95 °C block temperature and 105 °C lid temperature in a PCR cycler, cooled down at room temperature and stored on ice until usage. To form RNP complexes, gRNAs were pooled (final concentration 20 µM each) and mixed with the Cas9 enzyme (final concentration 1.2 µg/ml; TrueCut Cas9 Protein v2; Thermo Fisher Scientific) in IDTE buffer and incubated for 20 min at room temperature. Transfection was carried with

Nucleofection out using the SF Cell Line 4D-Nucleofector® X Kit (Lonza). 0.2 x $10^6$ were used per nucleofection well. Cells were harvested, washed with calcium-free PBS, and resuspended in nucleofection buffer (prepared according to manufacturers instructions. RNP-complex was added in a ratio of 1:3, followed by an incubation at room temperature for 2 min. Nucleofection was carried out with the 4D-Nucleofactor X Unit applying the program FF-120. Immediately afterwards, cells were transferred to a 24 well plate containing 1 ml pre-warmed culture medium. To generate single cell clones, a limiting dilution was performed in the following passage.

*Co-culture of PCI tumor spheroids and human immune cells*

MNCs for co-culture with PCI-15 spheroids were isolated via density gradient centrifugation, and granulocytes by hypotonic lysis. For addition of unstimulated immune cells (first day of co-culture), granulocytes and MNCs were mixed, and 0.1 x $10^6$ cells added to the spheroids. For pre-stimulation of T cells, fraction of T cells in MNCs was determined according to cell size using the CASY Cell Counter, and anti-CD3/CD28 Dynabeads (Thermo Fisher Scientific= in a bead-to-cell ratio of 1:2 were added to the MNCs. Next day, beads were removed, and 0.1 x $10^6$ T cells added to the co-culture for further 48 h. 72 h after starting co-culture, non-infiltrated immune cells were washed away, and co-cultures were further processed for flow cytometry analyses, Live Cell Imaging or embedded for IHC and DESI-MRM-MSI.

Analysis of spheroid lysis was performed using the Incucyte Live Cell Imaging system (Sartorius). Non-infiltrated immune cells were carefully removed by washing and spheroids transferred to a fresh Poly-Hema coated culture plate maininting the respective treatment (end volume 200 µl). To monitor cell viability, 1 µl Cyto3D® Live-Dead Assay Kit (The Well Bioscience) per well was added.

For IHC analysis, at least 10 spheroids per condition were embedded. If metabolic imaging was performed, consecutive sections were used for IHC. Tumor spheroids were placed on the stamp of a 1 mL syringe, embedded in hydrogel matrix and placed into embedding molds. The bottom of the embedding mold was dipped in liquid nitrogen until the matrix was completely frozen. Molds were covered with parafilm and stored at -80 °C until being processed. For IHC, per experiment, multiple slides were examined to guarantee analyses of equal levels.

*Ex vivo tumor fragment culture*

Tissue was harvested, immediately transferred to the laboratory on ice and divided into fragments of 15 – 20 mg. Tumor fragments were minced (roughly 2 mm size) and cultivated in 2 mL RPMI 1640, penicillin (50 U/mL), streptomycin (50 µg/mL) and 10 % autologous serum in 12-well suspension culture plates with or without addition of 2 mM L-glutamine, 10 µM BPTES, 0.2 mM diclofenac (Sigma-Aldrich), 1 µM AZD3965 (AZD1 = MCT1 inhibitor), 1 µM AZD0095 (AZD4 = MCT4 inhibitor; both Hycultec), 20 mM L-glutamate (Sigma-Aldrich), 4 mM GSH-MEE (Sigma-Aldrich; dissolved freshly), 15 mM lactic acid (Sigma Aldrich) and 10 µg/mL anti-PD1 (BioXCell). For cultivation with different glucose concentrations, glucose-free RPMI1640 (PAN Biotech) was used and D-glucose (Sigma-Aldrich) was added at the respective concentrations. Immune cells were isolated from fragment tissue for flow cytometry as described above. For analysis of intracellular cytokine expression, fragments were treated with monensin for 2 h. Except for those fragments displayed in the comparison of immune infiltration and activity before and after culture, fragments were beforehand treated with phorbole-myristate-13-acetate (PMA, 0.018 µg/ml, Calbiochem, Merck Millipore) and Ionomycin (0.89 µM; Enzo life sciences) to induce maximal cytokine response. Digestion time was reduced to 1 h. For immunohistochemical analysis, fragments were carefully picked up

using a 1 ml pipette and transferred to fresh culture plate containing 1 ml cooled PBS. The bottom of an embedding mold was covered with hydrogel and placed on dry ice until freezing process started. The PBS-washed fragment was placed into the hydrogel, covered and the mold dipped into liquid nitrogen until the sample was entirely frozen. Processing for immunohistochemistry was performed as described.
Culture supernatants were stored at -80 °C for later analysis of metabolite and cytokine concentrations.

*Flow cytometry*

*Surface staining protocol*

0.5-1 x $10^6$ cells/tube were administered for staining of surface markers. 10 µl FcR-blocking reagent (Miltenyi Biotec) was administered for 10 min at 4 °C after washing with FACS buffer. Cells were stained with 80 µL Zombie-NIR (diluted 1:650 in PBS) or Zombie-Aqua (diluted 1:200 in PBS; both BioLegend) for 10 min at room temperature in the dark for live-dead discrimination. Subsequently, surface staining was performed in 50 µL Brilliant Stain Buffer (BD Biosciences) for 20 min at 4 °C. Antibodies are listed below.

*Intracellular staining of cytokines*

Intracellular staining for cytokines and Ki-67was performed in monensin-treated cells after 20 min fixation/permeabilization using the eBioscience™ Foxp3/Transcription Factor Staining Buffer (eBioscience, Thermo Fisher Scientific) according to the manufacturer's protocol. Staining was performed in FACS buffer for 20 min at 4 °C.

*Determination of cellular ROS levels*

Cellular ROS was determined by staining with 2′,7′-dichlorofluorescin diacetate ($H_2$DCFDA, Sigma-Aldrich). After surface staining, cells were resuspended in 500 µL FACS buffer. Staining was performed by addition of 10 µM $H_2$DCFDA, followed by incubation in the cell culture incubator for 20 min with capped tubes. Cells were washed with 3 mL ice-cold PBS and measurement was performed immediately.

*Western Blot Analysis*

For analysis of glutamine synthetase expression, PCI-15 cells were cultivated with and without 2 mM L-glutamine for 24 h, all other analysis were performed without prior treatment.Cells were washed 3x times with PBS, suspended in Cell lysis Buffer (Cell Signaling Technology), and stored at -80 °C. Samples were separated by 12 % SDS-PAGE. After transfer of proteins, PVDF membranes were blocked with 5 % skimmed milk (Sigma-Aldrich) in TBS buffer (supplemented with 0.1 % Tween-20) for 1 h. Incubation with primary antibodies was conducted at 4 °C overnight. Next day, membranes were incubated with the secondary antibody for 1 h at room temperature. For detection multiple proteins, membranes were stripped using the ReBlot Plus Strong Stripping Solution (Merck Millipore, Sigma-Aldrich). Detection was performed by chemiluminescence (ECL; Amersham Bioscience) and analyzed using the chemiluminescence system Fusion Pulse 6 (Vilber Lourmat).

### Information on antibodies

| Antigen | Source | Identifier |
|---|---|---|
| **Antibodies for Flow cytometry** | | |
| CCR7 (clone G043H7) | BioLegend | Cat # 353232 |
| CD11b (clone ICRF44) | BioLegend | Cat # 557918 |
| CD14 (clone 63D3) | BioLegend | Cat # 367140<br>Cat # 367114 |
| CD15 (clone W6D3) | BioLegend | Cat # 323040 |
| CD19 (clone SJ25C1) | BioLegend | Cat # 363020 |
| CD226 (clone 11A8) | BioLegend | Cat # 338334 |
| CD25 (clone S20019) | BioLegend | Cat # 385210 |
| CD3 (clone UCHT1) | BD Biosciences | Cat # 612940 |
| CD4 (clone SK3) | BioLegend | Cat # 344612 |
| CD45 (clone 2D1) | BioLegend | Cat # 368506 |
| CD45RO (clone UCHL1) | BioLegend | Cat # 304232 |
| CD56 (clone 5.1H11) | BioLegend | Cat # 362532 |
| CD8 (clone RPA-T8) | BD Biosciences | Cat # 563823 |
| CXCR3 (clone G025H7) | BioLegend | Cat # 353716 |
| CXCR4 (clone 12G5) | BD Biosciences | Cat # 563924 |
| CXCR6 (clone K041E5) | BioLegend | Cat # 356006 |
| Granzyme B (clone QA16A02) | BioLegend | Cat # 372206 |
| HLA-ABC (clone G46-2.6) | BD Biosciences | Cat # 565334 |
| HLA-DR (clone L243) | BioLegend | Cat # 307640 |
| IFNγ (clone 4S.B3) | BioLegend | Cat # 502560 |
| IFNγ (clone B27) | BD Biosciences | Cat # 563563 |
| Ki67 (clone Ki-67) | BioLegend | Cat # 350514 |
| PD-1 (clone Eh12.2H7) | BioLegend | Cat # 329908 |
| PD-L1 (clone MIH1) | BD Biosciences | Cat # 558065 |
| Perforin (clone dG9) | BioLegend | Cat # 308120 |
| TNF (clone Mab11) | BioLegend | Cat # 502936 |
| **Antibodies for IHC** | | |
| CD3 | Agilent | Cat # A045229-2 |
| TUNEL in situ Apoptosis Kit | Biomol | Cat # E-CK-A331.50 |
| **Antibodies for Western Blot** | | |
| Glutamine synthetase | Thermo Fisher Scientific | Cat # MA5-54014 |
| Glutaminase 1 | Cell Signaling Technology | Cat # 56750T |
| Aktin | Sigma-Adrich | Cat # A2066 |
| Goat anti-rabbit HRP | Agilent | Cat # P044801-2 |
| **Antibodies for Cell culture** | | |
| PD-1 (J116, blocking antibody) | BioXCell | Cat # BE0188 |
